# Structural Determinants of Redox Conduction Favor Robustness over Tunability in Microbial Cytochrome Nanowires

**DOI:** 10.1101/2023.01.21.525004

**Authors:** Matthew J. Guberman-Pfeffer

## Abstract

Helical homopolymers of multiheme cytochromes catalyze biogeochemically significant electron transfers with a reported 10^3^-fold variation in conductivity. Herein, classical molecular dynamics and hybrid quantum/classical molecular mechanics are used to elucidate the structural determinants of the redox potentials and conductivities of the tetra-, hexa-, and octaheme outer-membrane cytochromes E, S, and Z, respectively, from *Geobacter sulfurreducens*. Second-sphere electrostatic interactions acting on minimally polarized heme centers are found to regulate redox potentials over a computed 0.5-V range. However, the energetics of redox conduction are largely robust to the structural diversity: Single-step electronic couplings (⟨H_mn_⟩), reaction free energies 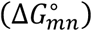, and reorganization energies (λ_mn_) are always respectively <|0.026|, <|0.26|, and between 0.5 – 1.0 eV. With these conserved parameter ranges, redox conductivity differed by less than a factor of 10 among the ‘nanowires’ and is sufficient to meet the demands of cellular respiration if 10^2^ – 10^3^ ‘nanowires’ are expressed. The ‘nanowires’ are proposed to be differentiated by the protein packaging to interface with a great variety of environments, and not by conductivity, because the rate-limiting electron transfers are elsewhere in the respiratory process. Conducting-probe atomic force microscopy measurements that find conductivities 10^3^-10^6^-fold more than cellular demands are suggested to report on functionality that is either not used or not accessible under physiological conditions. The experimentally measured difference in conductivity between Omc- S and Z is suggested to not be an intrinsic feature of the CryoEM-resolved structures.

---

Cytochrome ‘nanowires’ are polymeric multi-heme cytochromes that catalyze the transfer of electrons across microbe-microbe^1^ and microbe-mineral^2–5^ interfaces. As terminal reductases in a widespread respiratory strategy^6,7^ of biogeochemical significance,^2–5^ and promising materials for bioelectronic technologies,^8–15^ cytochrome ‘nanowires’ have attracted much recent interest.

The known structures^16–18^ invariably feature a spiraling chain of (typically) alternating slipped and T-stacked bis-histidine-ligated *c*-type heme cofactors encased by a protein sheath (Figure 1). How differences within this architectural blueprint encode a 10^3^-fold variation in electrical conductivity^19^ is currently not well understood. Conductivity in cytochrome ‘nanowires,’ particularly under physiological conditions (not necessarily those of atomic force microscopy experiments), has previously been modeled as a cascade of reduction-oxidation (redox) reactions,^20–23^ but the structural determinants of the underlying redox potentials have not yet been elucidated for any of the known structures. These potentials can be tuned over a 0.8-V range by the protein matrix.^24^ Do selective pressures operate on the protein matrix to tune the redox potentials and thereby optimize the electrical conductivity of the chain of heme cofactors? These questions motivated the present Letter.

**Figure 1.**
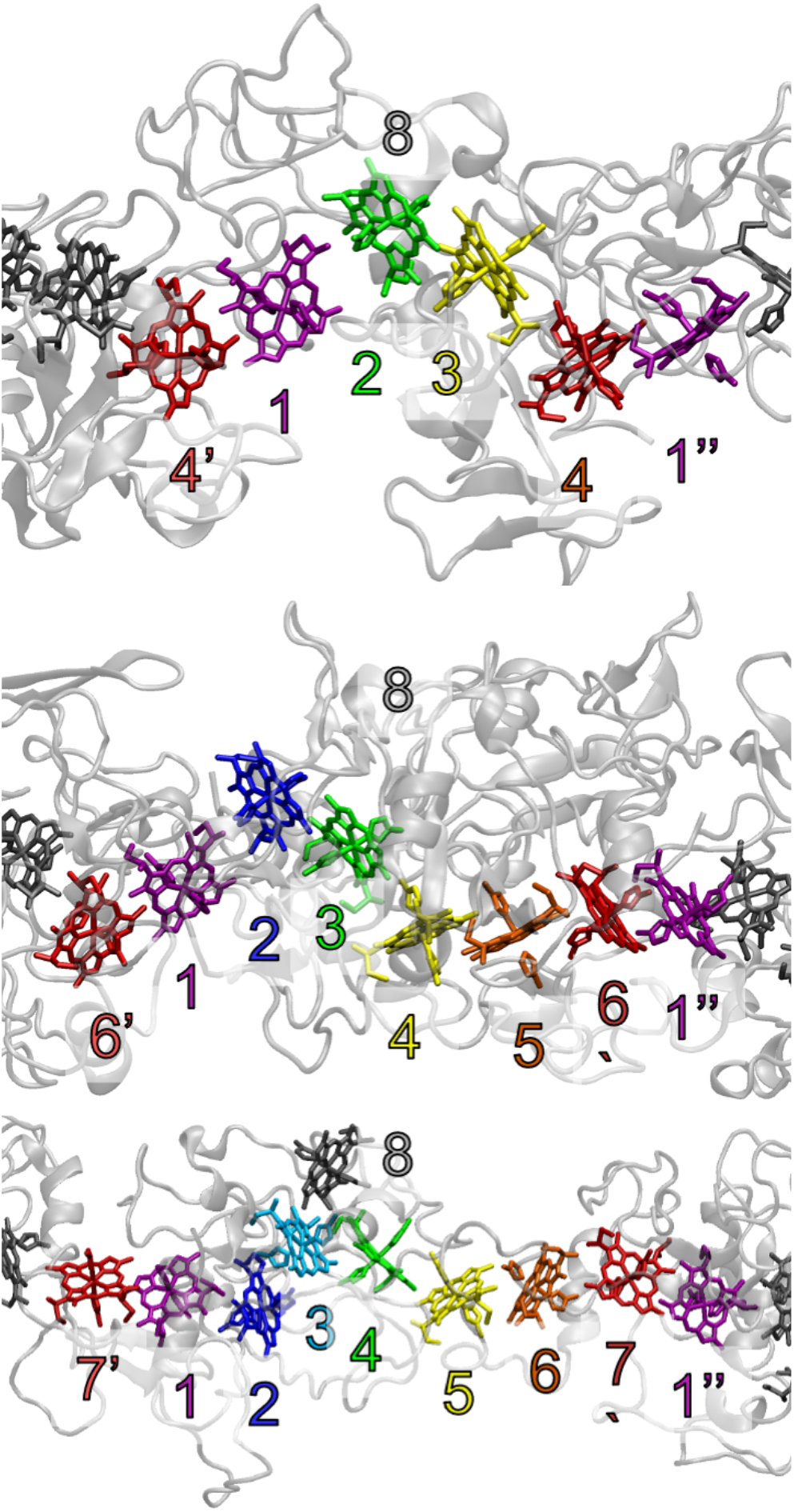
Structures of Omc- E (PDB 7TF6), S (PDB 6EF8), and Z (PDB 7QL5) from top to bottom, respectively, with the color and naming conventions used for the hemes in the present work indicated. Context will make it clear which set of hemes is being referenced. The figures were prepared using visual molecular dynamics (VMD) version 1.9.4a51.

Herein, second-sphere electrostatic interactions acting on largely non-polarized heme centers are shown to tune the redox potentials over a 0.5-V range in ‘nanowires’ of the tetra-, hexa-, and octaheme outer-membrane cytochromes (Omc-) E, S, and Z, respectively, from *Geobacter sulfurreducens*. These are the only cytochrome ‘nanowires’ so far with experimentally determined structures.^16–18^ Despite the variation in redox potentials, energetic parameters for electron transfer are found to reside within conserved parameter ranges, leading to the proposed evolutionary preference for functional robustness over tunability.

Inspired by prior work on the controlling factors for heme redox potentials,^25–29^ the redox potential of each heme in the hydrated ‘nanowires’ was approximated as a sum of conformational 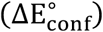 and electrostatic 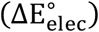 perturbations to the potential of the fully optimized heme cofactor in vacuum 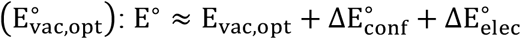.

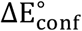 was assessed by transferring into the gas phase thermally averaged ensembles of heme conformations generated by classical molecular dynamics within the hydrated ‘nanowires.’ The *in vacuo* redox potentials were approximated as averages of vertical ionization potentials and vertical electron affinities from density functional theory computations on these heme conformers.

The lefthand side of Figure 2 shows that the conformations that were imposed by the Omc- E, S, and Z proteins (Tables S1-S6) minimally (< |0.09| V; Table S7-S9) shift E° relative to the fully optimized heme group (Tables S10, S11): 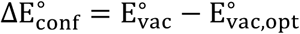. The total out-of-plane distortion (d_oop_) of the heme macrocycles is 1.9 – 5.7-times larger than the total in-plane (d_ip_) distortion. No simple relationship between d_oop_ and 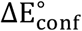 was found because the hemes exhibit a mixture of displacements along vibrational modes that share irreducible symmetry representations with different frontier molecular orbitals.^30,31^ Some hemes are principally ruffled (OmcE: #1 and #4; OmcS: #1, #2, #3 and #5; OmcZ: #1, #5, #6 and #7), principally saddled (OmcS: #4 and #6; OmcZ: #3 and #4), equally ruffled and saddled (OmcE: #3; OmcZ: #8), equally ruffled and waved (OmcZ: #2), or equally saddled, ruffled, and waved (OmcE: #2).^32^ Thus, little or no selection for a particular heme conformation seems to be conserved among the Omc- E, S, and Z ‘nanowires.’ This observation does not preclude the possibility that site-specific heme conformations may be conserved in homologous cytochromes.

**Figure 2.**
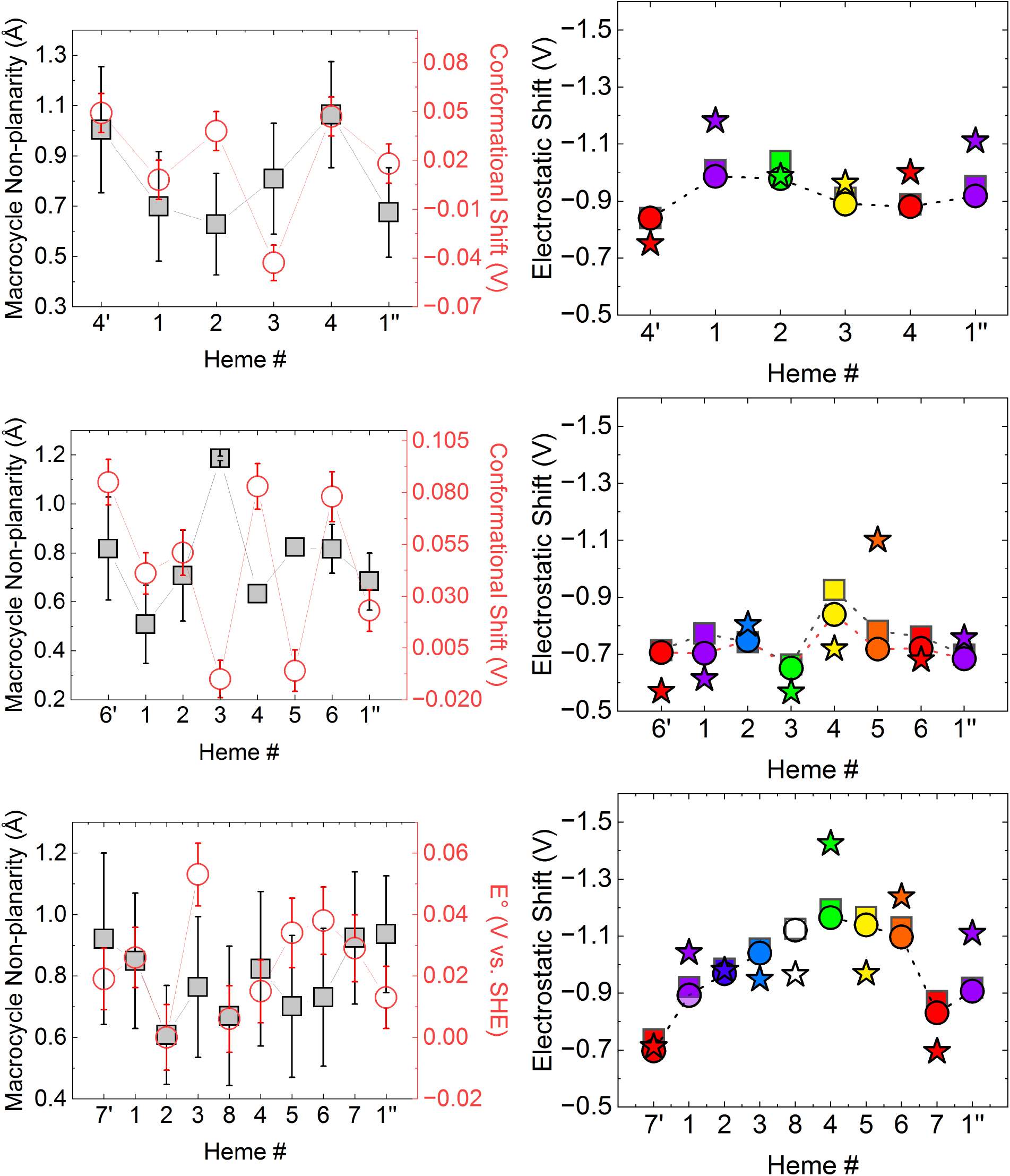
Analysis of (*left*) conformational and (*right*) electrostatic shifts imposed by the hydrated protein environment on the heme cofactors of Omc- E, S, and Z from top to bottom. Circles, squares, and stars on the righthand side indicate the electrostatic shift in redox potential approximated, respectively, as the difference in potentials between the hydrated protein and vacuum states, the sum of protein and solvent contributions in the hydrated protein, or the Coulombic energy for heme oxidation.

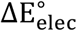 was estimated in three different ways (Eq. 1): (1) The difference between E° in the aqueously solvated protein 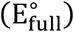 and in vacuum 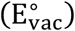, (2) the sum of protein – 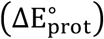 and solvent – 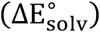 induced shifts to 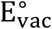, and (3) the change in Coulombic interaction energy (ΔE_coul_) for heme oxidation in the protein environment assessed with fixed (non-polarizable) atomic partial charges.

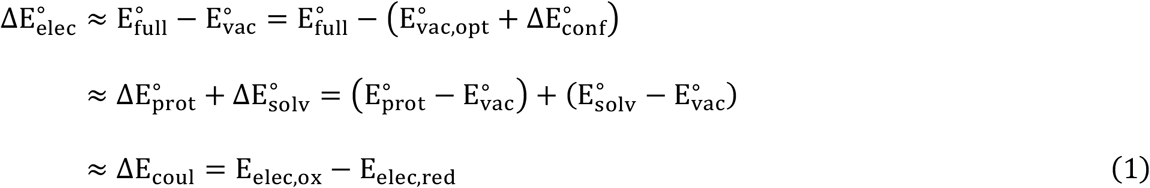

The righthand side of Figure 2 (Tables S7-S9) shows that all three estimates of 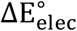 are in reasonable agreement and exceed 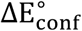 by roughly an order-of-magnitude: 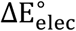 is −0.65 – −0.84 (OmcS), −0.88 – −0.99 (OmcE), or −0.70 – −1.17 (OmcZ) V versus <|0.09| V for 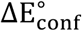. Thus, second-sphere electrostatics primarily tune heme redox potentials in these cytochrome ‘nanowires.’ Interestingly, the analysis showed 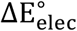 to result from a near complete cancellation of 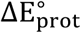 and 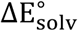: The excess negative charge of the protein and the excess positive charge of counterions in the surrounding solvent shift E° by nearly equal amounts in opposite directions.

To get an intuitive interpretation of 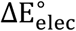, the hemes of Omc- E, S, and Z were transferred into solvents of different static dielectric constants (ϵ_s_)—other than 1.0 for vacuum—to find what homogeneous solvents reproduce the shifts in redox potentials exerted by the hydrated ‘nanowires.’

Increasing ϵ_s_ in Figure 3 from 1.0 to 1.9 (vacuum to *n*-hexane) captures 62 – 77% of the 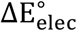 exerted by the binding sites of OmcS (Table S12). This large (−0.500 – −0.541 V) effect is understandable from the perspective of the Born equation for solvation of an ion: A non-polar solvent with ϵ_s_ = 2 provides half the solvation free energy as a perfect conductor with ϵ_s_ = ∞. Also, the hemes in OmcS are buried (solvent accessible surface area <70 Å^2^ of a maximum 882 Å^2^ for the heme group) in the protein, so the binding sites are expected to be largely non-polar in character. Note that the heme group is taken here to mean the bis-imidazole ligated heme macrocycle not including the propionic acid substituents.

**Figure 3.**
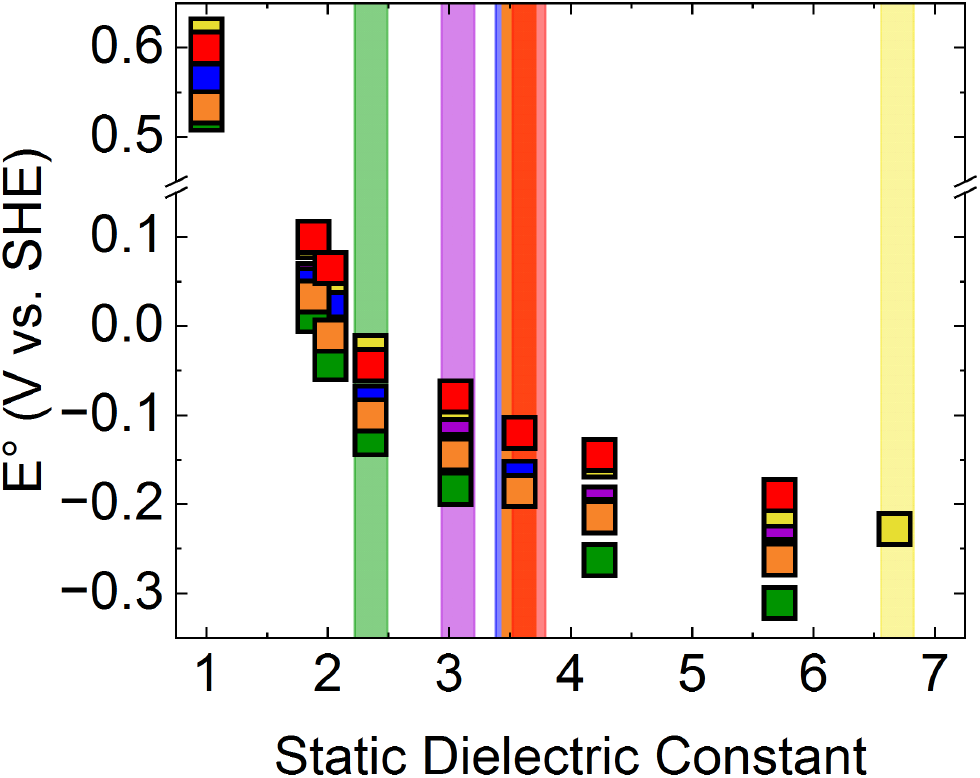
Solvent dependence of the heme redox potentials. The vertical strips indicate the effective static dielectric constant at which the color-coded hemes obtain the redox potential computed within the context of the hydrated OmcS protein.

As ϵ_s_ is further increased, the hemes titrate out to the E° found in OmcS at 2.35 (heme #3), 3.05 (Heme #1), 3.58 (Hemes #2, #5, #6), and 6.67 (Heme #4) (Figure 3, Table S12). Values of ϵ_s_ in the range of 3 to 7 are consistent with prior work characterizing the interior of proteins.^33,34^ Interestingly, hemes #3 and #4 experience dielectric extremes in the protein, even though these hemes are in van der Waals contact. The explanation for this fact requires a microscopic description of specific interactions, as developed below.

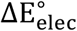 is larger in OmcE (−0.88 – −0.99 V) and OmcZ (0.70 – −1.17 V) than OmcS, (−0.65 – −0.84 V), suggesting even larger ϵ_s_ are needed to reproduce the influence of the protein environment. However, the effect of a homogeneous dielectric on heme redox potential saturates-out by ϵ_s_ ≈ 20 (*e.g*., acetone), in which a bis-imidazole-ligated model heme group is predicted to have an E° of −0.363 V vs. SHE (Table S13). Increasing ϵ_s_ by an additional 88 units to that of formamide only increased E° by 0.023 V (Table S13). The failure to model the electrostatic influence of the heme binding sites of Omc- E and Z with an implicit homogeneous solvent underscores the importance of specific interactions with the heterogeneous hydrated-protein ‘solvent.’ Identifying these interactions is possible by finding a way to decompose 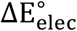 for the full environment into contributing structural features.

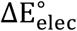 was computed from wavefunctions for the heme cofactor polarized by the electrostatic environment. By contrast, ΔE_coul_, which reproduces 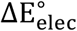 with a mean-unsigned error (MUE) of <0.16 eV, was computed from classical electrostatics using fixed atomic partial charges. Note that the MUE for the approximate density functional (B3LYP) used to compute 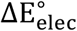 was reported as ~0.11 V with respect to *ab initio* theory for related systems.^35^ That 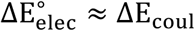 (Figure 2) suggests polarizability of the heme centers is a minor factor. In fact, redox potentials evaluated in the protein context with either unrelaxed (frozen vacuum-optimized) or relaxed (environmentally polarized) heme electron densities agreed within thermal energy for a one-electron process at 298 K (Tables S14-S16). This is an intriguing result given the prior finding that polarizability plays a role in lowering the activation energy for electron transfer from the differently-ligated heme cofactor of cytochrome *c*.^36,37^

Given that 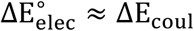 the factors controlling E° should be decomposable into pairwise additive contributions. Energy decomposition schemes are in general not uniquely defined, but two chemically motivated approaches involve partitioning interacting groups by physicochemical character (e.g., non-polar, aromatic, polar, acidic, basic, etc.) (Tables S17-S19), or by amino acid residue (Tables S20-S22).

Testing the latter approach, 11 of the largest per-residue contributions to the change in interaction energy upon oxidation of hemes #3 and #4 in OmcS were compared to the change in redox potentials produced by switching on/off electrostatic interactions with these residues (Tables S23-S29). Per-residue 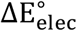 and ΔE_coul_ correlated with an R^2^ of 0.75 and 0.89 for hemes #4 and #3, respectively, and 0.71 overall. The correlation fell to an R^2^ of 0.43 for heme #5, which was included as a negative control: The examined residues were not among the strongest interactions for this heme and the energetic shifts were close to the noise from thermal sampling.

The per-residue analysis revealed that Arg-187 and Arg-333 are principally responsible for taking heme #3 from having one of the most negative potentials in vacuum because of its protein-imposed conformation to one of the most positive potentials in OmcS. By contrast, no single-dominant interaction sets the redox potential of heme #4 in OmcS.

Generalizing the analysis to the other ‘nanowires’, 11 – 27 positive and 11 – 27 negative interactions make contributions of >|0.025| V to the redox potential of every heme in Omc- E, S, and Z. Redox potentials reflect an incomplete cancellation of many competing factors. This insight is a long-established principle,^27^ but deserves emphasis here to inject humility into discussions of cytochrome ‘nanowires’ as tunable or programmable biomaterials.

The sum of negative per-residue contributions to the redox potentials always exceeded the sum of positive per-residue contributions. The excess negative electrostatic interaction, averaged over all the hemes in each ‘nanowire’, became more negative (−0.81 → −1.06 → −1.13 eV) in parallel with a lowering of the vertical electron affinity from the averaged vacuum value by −1.49, −1.75, and −2.02 eV from Omc- S to E to Z. Thus, these cytochrome ‘nanowires’ achieve different redox potential ranges and macroscopic midpoints by differently stabilizing the oxidized state of the bound hemes.

One specific interaction responsible for this effect that is unique to OmcZ and not captured by a dielectric solvent model for the hydrated protein is an axial histidine ligand-to-propionate heme sidechain H-bond on the same or different cofactor (Figure 4). H-bonds of this sort position the negatively charged propionates very close to the heme macrocycle, and thereby contribute significantly to the more negative E°s found in Omc- Z (Table S9) versus E (Table S7) or S (Table S8). Three of these H-bonds are conserved in the fully oxidized and single-heme reduced states, whereas three more are redox-state dependent (Figure 4; Table S30). Five of the interactions form an inter-heme H-bonding network that may facilitate electron transfer, or proton-coupled electron transfer in the ‘nanowire.’

**Figure 4.**
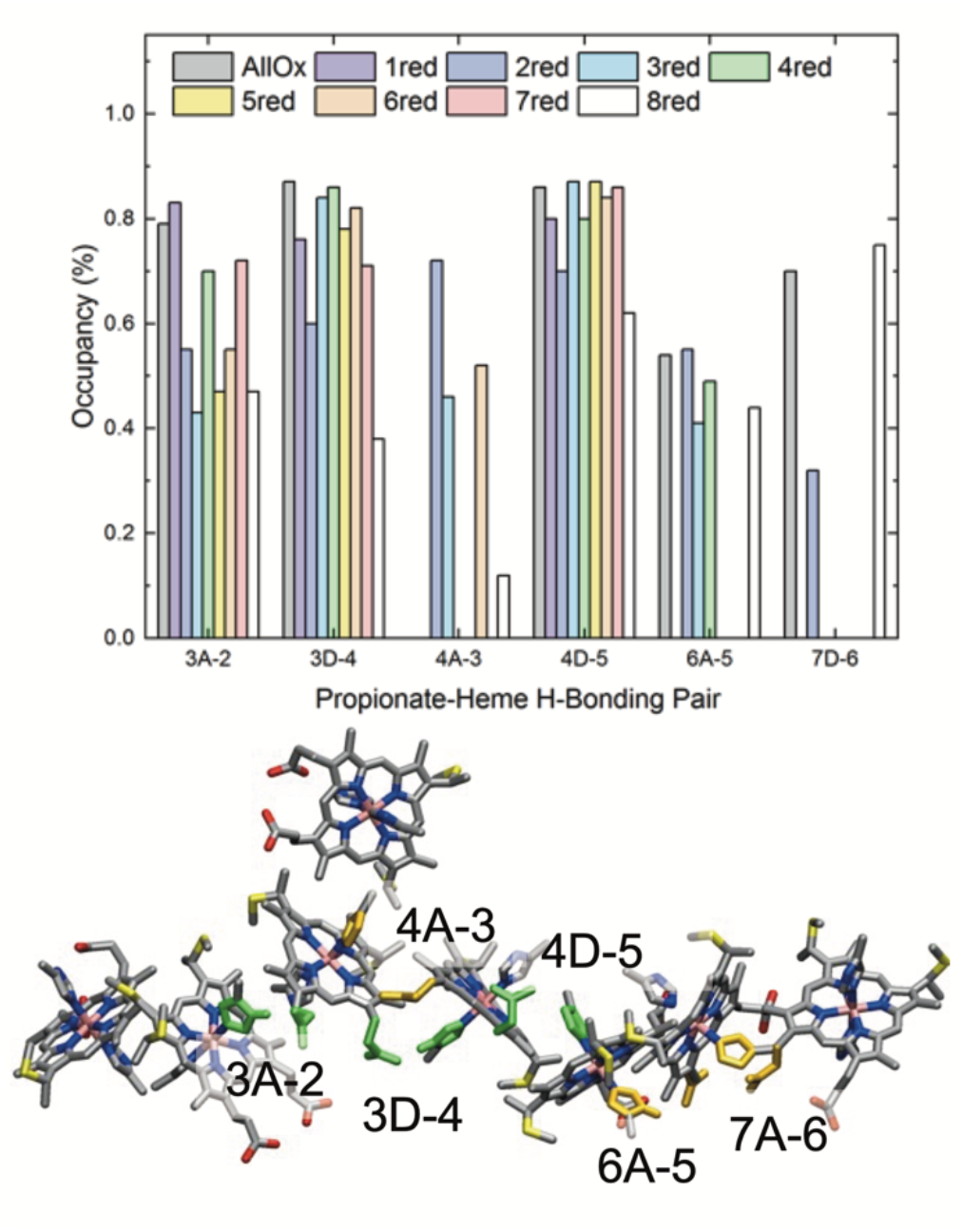
(top) Assembly of hemes in OmcZ with H-bonded axial histidine ligands and propionates colored green and yellow if the interactions are redox-state independent or dependent, respectively. (bottom) Occupancy of the indicated H-bonds.

To summarize so far, the approximation 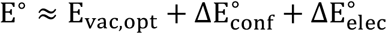 simplifies to E^°′^ ≈ E_coul_, where the constant E_vac,opt_ has been subsumed into E°, ΔE_conf_ has been dropped because it is an order-of-magnitude smaller than 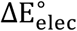 and 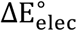 has been replaced with ΔE_coul_ because polarizability is a relatively minor secondary effect. ΔE_coul_ can be partitioned into additive pairwise interactions to identify the most important structural determinants setting E^°′^. This conclusion lays the groundwork for a (prudently exercised) *in silico* mutagenesis strategy (Figure S1) for specific applications.

The relationship E^°′^ ≈ ΔE_coul_ gives insight into how evolution and rational design efforts may modulate redox conductivity in cytochrome ‘nanowires’: If changes in the electrostatic environment differentially shift the E^°′^ of some heme groups, the ΔE^°′^ between adjacent hemes can change, and with it, the free energy (ΔG°) for electron transfer, which enters the exponential of the Marcus-theory^38,39^ rate expression (Eq. 2).

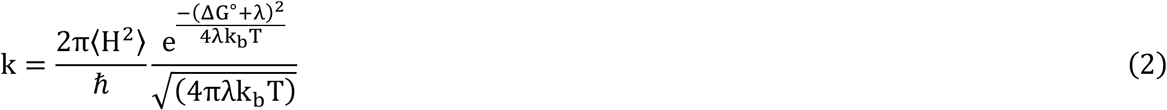

The other energetic parameters in Eq. 2 are the reaction reorganization energy (λ_rxn_) and the donor-acceptor electronic coupling (⟨H⟩). k_b_, T, and ℏ, signify, respectively, the Boltzmann constant, absolute temperature, and the Plank constant.

Has evolution used this redox potential-tuning strategy to modulate conductivity in cytochrome ‘nanowires’ by a reported^19^ 10^3^-fold? Very unlikely seems the answer: Despite the 0.5-V range in computed E° for Omc- E, S, and Z, ΔG° is always <|0.26| eV (Figure 5, top panel; Tables S31-S33).

**Figure 5.**
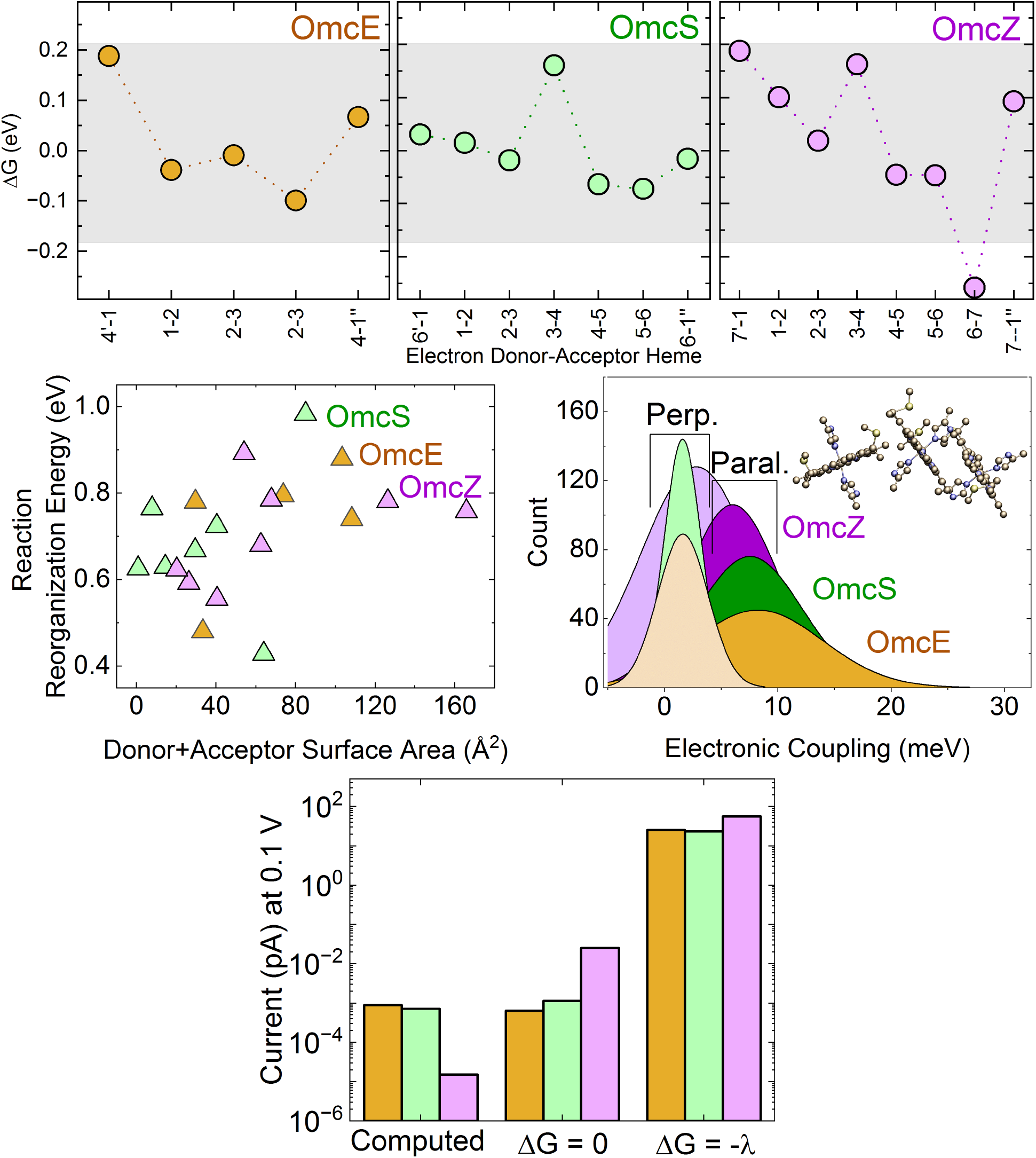
Comparison of the reaction (ΔG°) and reorganization (λ_rxn_) free energies and electronic couplings (⟨H⟩) for Omc-E, S, and Z, and the predicted currents.

Moreover, in the hydrated state, ΔG° is typically a minor component of the electron transfer activation barrier 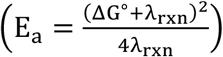 for these systems compared to *λ*_rxn_, which is 0.5 – 1.0 eV. As expected, there is some relationship between λ_rxn_ and the solvent accessibility of the heme macrocycles (Figure 5, middle left);^40^ both of these parameters being similar across the three ‘nanowires.’ λ_rxn_ is also more than 20-times larger than the ⟨H⟩s, which are sub-0.026 eV in Omc- E, S, and Z (Figure 5, middle right). The T-stacked hemes in OmcZ tend to be ~1 Å closer than in Omc- E or S (Table S34-36), but the geometrical difference has a relatively minor influence on the thermal distribution of ⟨H⟩ (Figure 5, middle right) and cannot by itself explain the 10^3^-fold higher conductivity. The result that λ_rxn_ ≫ ⟨H⟩ justifies the use of the non-adiabatic form of the Marcus rate expression in Eq. 2 to compute redox conductivity.

Within the conserved parameter ranges for ΔG°, λ_rxn_, and ⟨H⟩, the computed redox current for a 300 nm filament under low (0.1 V) applied bias is predicted to increase in the order Omc- Z (1.5 × 10^−2^) < S (7.2 × 10^−1^) < E (8.9 × 10^−1^ fA) (Figure 5, bottom). Compared to the 10^2^-fA current discharged by an entire *Geobacter sulfurreducens* cell,^41,42^ 6667, 138, or 115 ‘nanowires’ of exclusively the Omc- Z, S, or E variety would be needed. Note that the 5-fold smaller conductivity predicted for OmcZ is hardly significant: The computed lower conductivity is attributable to a strongly exergonic (ΔG° = −0.250 eV) heme #6 → #7 electron transfer step that creates a trap state. If the ΔG° for this step is overestimated by −0.050 eV, which is comparable to the standard-error-of-the-mean for this quantity, the computed current would be 10-times larger, and only 641 OmcZ ‘nanowires’ would be needed exclusively for cellular respiration. In fact, in the limit of all free-energy optimized rates, Omc- E, S, and Z have essentially the same conductivity, supporting currents of 20-60 fA at 0.1 V (Table S38).

The expression of hundreds-to-thousands of ‘nanowires’ by a single cell seems biologically reasonable to the present author. The prediction of functional similarity among all the known ‘nanowires’ also seems sensible given the expectation that catalytic centers, not redox cofactor chains, form the bottlenecks in biology.^43,44^ Formation of the pre-cursor complex between a ‘nanowire’ and molecular electron donors and acceptors (*i.e*., collisional diffusion, molecular recognition, and protein-protein binding) likely makes the injection or ejection of charges into the ‘nanowires’ slower than the movement of charge through a densely packed and rigid array of redox centers.

If ‘nanowires’ are not the rate-limiting step for extracellular electron transfer, little evolutionary pressure can be expected to differentiate them by conductivity. Instead, evolution will act to conserve the central heme architecture—not because it is optimal but because it is sufficient—and work to adapt the protein sheath to the particularities of the habitat for the microorganism. It is advantageous for the conductivity of the heme chain to be robust to rapid or large-scale mutational alterations to the surrounding protein so the same ‘solution’ to the respiratory problem can be used over-and-over again. This is a general theme in biology: Conserve the functional entity and simply adapt its interface or packaging for a great diversity of circumstances (e.g., light harvesting antenna complexes that allow photosynthetic organisms with the same photosystems to inhabit very different spectral niches.)

In the context of Omc- S and Z, *G. sulfurreducens* likely switches to overexpress OmcZ at an electrode^19^ not because the heme chain of OmcZ is intrinsically much more conductive, but because the greater aggregation propensity and exposure of the heme groups more readily facilitates inter-protein electron transfer than is possible with OmcS to build a conductive biofilm matrix.

From the perspective of conducting-probe atomic force microscopy (CP-AFM) measurements,^16,19^ however, the computed conductivities are too small by 10^4^- and 10^9^-fold for Omc- S and Z respectively; no data is currently available for OmcE. That is, CP-AFM experiments indicate that a 300-nm cytochrome ‘nanowire’ under a 0.1-V bias carries a thousand- to a million-fold greater current than produced by an entire *G. sulfurreducens* cell. The large discrepancy between cellular physiology and CP-AFM measurements suggests that either the Omc- S and Z ‘nanowires’ have a tremendous current-carrying capacity that is largely unused by a *G. sulfurreducens* cell, or the CP-AFM measurements are reporting on a biologically irrelevant conduction mechanism. Note that *G. sulfurreducens* will not exploit excess available potential energy if an anode is set at a potential more than 0.1-V above the thermodynamic limit for extracellular electron transfer.^45^

It is not surprising that the computations on OmcZ do not uphold the reported greater conductivity by CP-AFM, because the cryogenic electron microscopy (CryoEM) structure used as input to the computations disagrees with the other characterization details presented in that report, as already noted.^46^ Furthermore, it is very unlikely that the structures resolved by CryoEM and present in the CP-AFM measurements are the same, because in the latter, a relatively strong compressional force of 50 nN is applied to the protein.^47,48^ Given that OmcZ is more flexible than OmcS, this compressional force is likely to differentially change the conformation of OmcZ, and that may contribute to the difference in measured conductivity between the two ‘nanowires.’ This is particularly the case since the length dependence of the current was measured with increasing distance from the Au electrode (see Fig.3 in Ref. 19), meaning that the current in subsequent measurements had to pass from the tip to the other electrode through previously compressed sections of the protein.

In closing, a mechanistic picture appliable to all known cytochrome ‘nanowires’ has been developed in which second-sphere electrostatic interactions acting on largely non-polarized heme centers regulate redox potentials. Despite variations in E°, potential *differences* for adjacent hemes fall within a conserved range for Omc- E, S, and Z, as do the other energetic parameters of non-adiabatic redox reactions. Thus, structural determinants of redox conductivity in cytochrome ‘nanowires’ favor functional robustness over tunability in response to, for example, variation in protein sequence, a finding consistent with the high mutational rate among multi-heme cytochromes.^17^

## Computational Methods

Theoretical background and technical details for the molecular dynamics (MD) and quantum mechanical/molecular mechanical (QM/MM) density functional theory (DFT) protocols used in this work were extensively described earlier.^23^ Here, focus is given to a high-level summary of the computations.

Trimeric assemblies of Omc- E (PDB 7TF6), S (PDB 6EF8), and Z (PDB 7QL5) were used as filament models. The filaments were assigned standard protonation states for pH 7 and immersed in aqueous solutions having sufficient Na^+^ ions to neutralize the net −3, −6, and −15*e* charge-per-subunit of Omc- E, S, and Z, respectively. Each system then underwent minimization, thermalization to 300 K, and density equilibration under a 1.0 bar atmosphere. Production-stage trajectories for each filament were propagated in the *NVT* ensemble for all-heme-oxidized and single-heme-reduced microstates. The reduced heme in the latter set of simulations was one of the cofactors in the central subunit of the filament model, the last heme of the preceding subunit, or the first heme of the proceeding subunit. In total, 7, 9, and 11 MD simulations with accumulated times of 2.4, 2.1, and 1.8 μs were conducted, respectively, for Omc- E, S, and Z. Additionally, the single-heme-reduced trajectories for OmcZ were repeated in methanol (1.8 μs) and chloroform (1.7 μs) to mimic the aggregated or membrane-associated state.

For each of these simulations, 141 (OmcE) or 180 (Omc- S and Z separately) frames were selected after at least 36-ns of the production simulations at a 200-ps interval. The vertical energy gaps—ionization potentials and electron affinities were computed for each heme conformer using B3LYP with a mixed double-ζ basis set (LANL2DZ for Fe and 6-31G(d) for H, C, N, and S). Redox potentials were approximated with linear response theory as the average of the vertical ionization potential and the negative of the vertical electron affinity, set relative to the absolute potential of the standard hydrogen electrode (SHE), and corrected for the integrated heat capacity and entropy of the electron according to the electron convention and Fermi-Dirac statistics.

H-bonding networks and solvent accessibility were examined, respectively, with CPPTRAJ^49,50^ of the AmberTools package and Visual Molecular Dynamics (VMD).

## ASSOCIATED CONTENT

### Supporting Information

The Supporting Information is available free of charge on the ACS Publications website.

Tables detailing the normal coordinate structure decomposition analysis of heme conformations, energetic partitioning of redox potentials, assessment of the influence of polarizability and dielectric media on redox potentials, inter-heme H-bonds. Marcus-theory electron transfer parameters and predicted redox currents, minimum inter-heme distances, electron transfer reorganization energies for OmcZ in methanol and chloroform, and the length of the performed molecular dynamics simulations. Also a figure showing the proposed Effective Charge on Last Common Atom approach to *in silico* mutagenesis.

## AUTHOR INFORMATION

### Author Contributions

The research described herein was initiated and conducted solely by the corresponding author. Acknowledgements are given below to others who provided insightful advice on related questions.

### Notes

The author declares no competing financial interest.

## ACKNOWLEDGMENT

This research was supported by the National Institute of General Medical Sciences of the National Institutes of Health under Award Number 1F32GM142247-01A1. All calculations were performed using resources at the Yale High Performance Computing Center. Implementations of the diffusion kinetic model used to compute redox currents was kindly provided by Fredrik Jansson. Gratitude is expressed to Prof. Nikhil S. Malvankar for advanced access to the cryogenic electron microscopy structure of OmcZ. I wish to deeply thank Michael Buehl and Jonathan Colburn for a useful discussion on the partitioning of redox potentials into per-residue contributions. I also wish to thank Prof. Dmitry Matyushov and Ssetare Mostajabi for brining the issue of heme polarizability to my attention.

