## Supporting Information for "Structural Determinants of Redox Conduction Favor Robustness over Tunability in Microbial Cytochrome Nanowires"

Table S1. In-plane normal coordinate structure decomposition for the hemes of OmcE using the minimal basis set of vibrational modes.

| Heme # | B2g | B1g | Eux | Euy | A1g | A2g | SSD | Err |
|---|---|---|---|---|---|---|---|---|
| All Hemes Oxidized State | | | | | | | | |
| 4' | -0.006 ± 0.100 | 0.027 ± 0.119 | -0.004 ± 0.070 | -0.026 ± 0.076 | 0.110 ± 0.096 | 0.016 ± 0.041 | 0.233 ± 0.073 | 0.010 ± 0.003 |
| 1 | -0.029 ± 0.097 | 0.042 ± 0.108 | 0.026 ± 0.066 | -0.028 ± 0.077 | 0.146 ± 0.087 | 0.007 ± 0.038 | 0.247 ± 0.069 | 0.009 ± 0.003 |
| 2 | -0.025 ± 0.096 | 0.040 ± 0.106 | 0.005 ± 0.066 | -0.018 ± 0.064 | 0.168 ± 0.087 | 0.016 ± 0.035 | 0.252 ± 0.071 | 0.009 ± 0.003 |
| 3 | -0.001 ± 0.108 | -0.012 ± 0.122 | 0.033 ± 0.074 | 0.011 ± 0.069 | 0.147 ± 0.081 | 0.011 ± 0.041 | 0.250 ± 0.070 | 0.009 ± 0.003 |
| 4 | -0.029 ± 0.098 | 0.023 ± 0.104 | -0.014 ± 0.077 | -0.008 ± 0.072 | 0.100 ± 0.106 | 0.020 ± 0.041 | 0.227 ± 0.068 | 0.011 ± 0.004 |
| 1" | -0.039 ± 0.092 | 0.044 ± 0.113 | 0.037 ± 0.071 | -0.002 ± 0.081 | 0.156 ± 0.084 | 0.007 ± 0.041 | 0.255 ± 0.074 | 0.009 ± 0.003 |
| Single-Heme Reduced State | | | | | | | | |
| 4' | -0.002 ± 0.108 | 0.025 ± 0.120 | 0.008 ± 0.063 | -0.012 ± 0.078 | 0.146 ± 0.091 | 0.010 ± 0.041 | 0.249 ± 0.079 | 0.018 ± 0.087 |
| 1 | -0.053 ± 0.102 | 0.040 ± 0.111 | 0.006 ± 0.067 | -0.017 ± 0.075 | 0.158 ± 0.078 | 0.007 ± 0.041 | 0.256 ± 0.070 | 0.009 ± 0.003 |
| 2 | -0.014 ± 0.100 | 0.009 ± 0.101 | -0.035 ± 0.072 | -0.001 ± 0.072 | 0.179 ± 0.077 | 0.010 ± 0.046 | 0.259 ± 0.068 | 0.009 ± 0.002 |
| 3 | 0.001 ± 0.095 | -0.037 ± 0.120 | 0.016 ± 0.069 | 0.006 ± 0.071 | 0.121 ± 0.086 | 0.017 ± 0.036 | 0.231 ± 0.072 | 0.009 ± 0.003 |
| 4 | -0.010 ± 0.097 | 0.017 ± 0.101 | -0.008 ± 0.079 | 0.001 ± 0.075 | 0.177 ± 0.081 | 0.012 ± 0.043 | 0.259 ± 0.065 | 0.009 ± 0.003 |
| 1" | -0.034 ± 0.104 | 0.056 ± 0.106 | 0.002 ± 0.074 | 0.011 ± 0.074 | 0.164 ± 0.093 | 0.001 ± 0.038 | 0.263 ± 0.071 | 0.009 ± 0.003 |

Table S2. Out-plane normal coordinate structure decomposition for the hemes of OmcE using the minimal basis set of vibrational modes.

| Heme # | B2u | B1u | A2u | Egx | Egy | A1u | SSD | Err |
|---|---|---|---|---|---|---|---|---|
| All Hemes Oxidized State | | | | | | | | |
| 4' | 0.345 | -0.839 | 0.154 | 0.009 | -0.137 | 0.103 | 1.004 | 0.022 |
| | ± 0.254 | ± 0.259 | ± 0.131 | ± 0.142 | ± 0.138 | ± 0.091 | ± 0.251 | ± 0.010 |
| 1 | -0.017 | -0.485 | 0.098 | -0.210 | -0.041 | 0.066 | 0.699 | 0.024 |
| | ± 0.293 | ± 0.276 | ± 0.152 | ± 0.138 | ± 0.166 | ± 0.103 | ± 0.218 | ± 0.012 |
| 2 | 0.266 | -0.227 | -0.109 | -0.241 | 0.032 | 0.041 | 0.628 | 0.024 |
| | ± 0.297 | ± 0.268 | ± 0.151 | ± 0.162 | ± 0.150 | ± 0.096 | ± 0.202 | ± 0.012 |
| 3 | 0.563 | -0.414 | 0.149 | -0.109 | 0.151 | 0.027 | 0.809 | 0.025 |
| | ± 0.234 | ± 0.208 | ± 0.136 | ± 0.139 | ± 0.120 | ± 0.088 | ± 0.221 | ± 0.012 |
| 4 | 0.410 | -0.883 | 0.173 | 0.017 | -0.032 | 0.120 | 1.064 | 0.023 |
| | ± 0.255 | ± 0.222 | ± 0.129 | ± 0.156 | ± 0.145 | ± 0.090 | ± 0.211 | ± 0.010 |
| 1" | 0.024 | -0.316 | 0.141 | -0.388 | 0.081 | 0.055 | 0.675 | 0.027 |
| | ± 0.254 | ± 0.265 | ± 0.151 | ± 0.152 | ± 0.145 | ± 0.085 | ± 0.178 | ± 0.011 |
| Single-Heme Reduced State | | | | | | | | |
| 4' | -0.182 | -0.517 | 0.243 | 0.033 | -0.021 | 0.084 | 0.731 | 0.027 |
| | ± 0.274 | ± 0.236 | ± 0.141 | ± 0.159 | ± 0.140 | ± 0.090 | ± 0.189 | ± 0.041 |
| 1 | -0.152 | -0.509 | 0.243 | -0.131 | 0.033 | 0.037 | 0.749 | 0.024 |
| | ± 0.317 | ± 0.276 | ± 0.164 | ± 0.154 | ± 0.139 | ± 0.086 | ± 0.228 | ± 0.012 |
| 2 | -0.048 | -0.188 | -0.022 | -0.157 | 0.100 | 0.013 | 0.551 | 0.022 |
| | ± 0.351 | ± 0.251 | ± 0.147 | ± 0.154 | ± 0.156 | ± 0.093 | ± 0.180 | ± 0.010 |
| 3 | 0.616 | -0.581 | 0.199 | -0.086 | 0.160 | -0.025 | 0.955 | 0.023 |
| | ± 0.243 | ± 0.214 | ± 0.144 | ± 0.143 | ± 0.137 | ± 0.086 | ± 0.222 | ± 0.010 |
| 4 | -0.036 | -0.549 | 0.247 | 0.134 | 0.054 | 0.018 | 0.748 | 0.022 |
| | ± 0.279 | ± 0.248 | ± 0.134 | ± 0.164 | ± 0.140 | ± 0.100 | ± 0.193 | ± 0.009 |
| 1" | -0.251 | -0.308 | 0.146 | -0.224 | 0.095 | 0.051 | 0.688 | 0.025 |
| | ± 0.316 | ± 0.290 | ± 0.166 | ± 0.167 | ± 0.169 | ± 0.097 | ± 0.208 | ± 0.013 |

Table S3. In-plane normal coordinate structure decomposition for the hemes of OmcS using the minimal basis set of vibrational modes.

| Heme # | B2g | B1g | Eux | Euy | A1g | A2g | SSD | Err |
|---|---|---|---|---|---|---|---|---|
| All Hemes Oxidized State | | | | | | | | |
| 6’ | -0.018 | 0.068 | 0.015 | -0.001 | 0.110 | 0.002 | 0.300 | 0.430 |
| | ± 0.108 | ± 0.115 | ± 0.069 | ± 0.070 | ± 0.318 | ± 0.039 | ± 0.253 | ± 2.319 |
| 1 | 0.001 | 0.041 | 0.013 | -0.001 | 0.184 | 0.017 | 0.265 | 0.008 |
| | ± 0.098 | ± 0.105 | ± 0.073 | ± 0.074 | ± 0.085 | ± 0.040 | ± 0.078 | ± 0.003 |
| 2 | -0.023 | 0.038 | -0.000 | -0.005 | 0.151 | 0.007 | 0.249 | 0.009 |
| | ± 0.107 | ± 0.104 | ± 0.072 | ± 0.075 | ± 0.083 | ± 0.043 | ± 0.067 | ± 0.003 |
| 3 | -0.011 | 0.026 | 0.032 | -0.011 | 0.053 | 0.007 | 0.208 | 0.013 |
| | ± 0.104 | ± 0.102 | ± 0.078 | ± 0.068 | ± 0.095 | ± 0.042 | ± 0.063 | ± 0.004 |
| 4 | -0.012 | 0.038 | 0.026 | 0.023 | 0.197 | 0.008 | 0.275 | 0.008 |
| | ± 0.096 | ± 0.110 | ± 0.069 | ± 0.069 | ± 0.076 | ± 0.044 | ± 0.065 | ± 0.003 |
| 5 | -0.016 | 0.006 | 0.007 | 0.007 | 0.120 | 0.006 | 0.224 | 0.009 |
| | ± 0.095 | ± 0.113 | ± 0.069 | ± 0.069 | ± 0.084 | ± 0.041 | ± 0.067 | ± 0.003 |
| 6 | 0.002 | 0.052 | 0.031 | 0.001 | 0.158±0 | 0.017 | 0.255 | 0.008 |
| | ± 0.100 | ± 0.117 | ± 0.069 | ± 0.073 | .083 | ± 0.036 | ± 0.075 | ± 0.003 |
| 1’’ | 0.016 | 0.031 | 0.011 | 0.030 | 0.163±0 | 0.015 | 0.255 | 0.009 |
| | ± 0.092 | ± 0.114 | ± 0.073 | ± 0.073 | .088 | ± 0.040 | ± 0.072 | ± 0.003 |
| Single-Heme Reduced State | | | | | | | | |
| 6’ | 0.001 | 0.045 | 0.008 | -0.001 | 0.165 | 0.016 | 0.254 | 0.009 |
| | ± 0.097 | ± 0.107 | ± 0.072 | ± 0.074 | ± 0.079 | ± 0.046 | ± 0.069 | ± 0.003 |
| 1 | 0.011 | 0.029 | 0.013 | 0.032 | 0.162 | 0.023 | 0.256 | 0.009 |
| | ± 0.095 | ± 0.113 | ± 0.078 | ± 0.071 | ± 0.076 | ± 0.042 | ± 0 .064 | ± 0.003 |
| 2 | -0.016 | 0.027 | -0.009 | -0.018 | 0.176 | 0.009 | 0.254 | 0.009 |
| | ± 0.093 | ± 0.105 | ± 0.063 | ± 0.073 | ± 0.080 | ± 0.038 | ± 0.069 | ± 0.003 |
| 3 | -0.005 | 0.019 | 0.043 | -0.012 | 0.064 | 0.011 | 0.209 | 0.013 |
| | ± 0.093 | ± 0.112 | ± 0.070 | ± 0.076 | ± 0.088 | ± 0.040 | ± 0.063 | ± 0.004 |
| 4 | -0.017 | 0.057 | 0.024 | 0.039 | 0.190 | 0.011 | 0.276 | 0.009 |
| | ± 0.098 | ± 0.109 | ± 0.066 | ± 0.073 | ± 0.079 | ± 0.039 | ± 0.066 | ± 0.003 |
| 5 | -0.026 | -0.006 | 0.010 | -0.025 | 0.130 | 0.000 | 0.239 | 0.009 |
| | ± 0.102 | ± 0.114 | ± 0.075 | ± 0.077 | ± 0.083 | ± 0.041 | ± 0.066 | ± 0.003 |
| 6 | 0.009 | 0.078 | 0.002 | 0.006 | 0.148 | 0.006 | 0.255 | 0.009 |
| | ± 0.093 | ± 0.108 | ± 0.065 | ± 0.079 | ± 0.098 | ± 0.043 | ± 0.072 | ± 0.003 |
| 1’’ | -0.001 | 0.045 | 0.008 | 0.031 | 0.154 | 0.021 | 0.248 | 0.009 |
| | ± 0.101 | ± 0.111 | ± 0.067 | ± 0.063 | ± 0.086 | ± 0.039 | ± 0.072 | ± 0.003 |

Table S4. Out-plane normal coordinate structure decomposition for the hemes of OmcS using the minimal basis set of vibrational modes.

| Heme # | B2u | B1u | A2u | Egx | Egy | A1u | SSD | Err |
|---|---|---|---|---|---|---|---|---|
| All Hemes Oxidized State | | | | | | | | |
| 6’ | -0.529 | -0.403 | 0.202 | -0.144 | -0.051 | 0.066 | 0.817 | 0.062 |
| | ± 0.273 | ± 0.231 | ± 0.141 | ± 0.163 | ± 0.137 | ± 0.090 | ± 0.210 | ± 0.290 |
| 1 | 0.042 | -0.269 | 0.038 | -0.158 | 0.093 | 0.005 | 0.508 | 0.023 |
| | ± 0.237 | ± 0.227 | ± 0.139 | ± 0.153 | ± 0.131 | ± 0.088 | ± 0.160 | ± 0.010 |
| 2 | -0.261 | -0.509 | 0.030 | -0.089 | 0.093 | 0.091 | 0.707 | 0.022 |
| | ± 0.235 | ± 0.237 | ± 0.134 | ± 0.150 | ± 0.143 | ± 0.099 | ± 0.185 | ± 0.009 |
| 3 | 0.048 | -1.136 | 0.061 | -0.122 | -0.005 | -0.076 | 1.186 | 0.021 |
| | ± 0.190 | ± 0.185 | ± 0.119 | ± 0.124 | ± 0.136 | ± 0.079 | ± 0.182 | ± 0.009 |
| 4 | -0.455 | -0.158 | 0.101 | -0.160 | -0.084 | 0.004 | 0.632 | 0.021 |
| | ± 0.219 | ± 0.238 | ± 0.120 | ± 0.127 | ± 0.125 | ± 0.084 | ± 0.182 | ± 0.010 |
| 5 | -0.237 | -0.658 | 0.005 | -0.109 | 0.194 | 0.004 | 0.823 | 0.025 |
| | ± 0.246 | ± 0.231 | ± 0.119 | ± 0.153 | ± 0.131 | ± 0.084 | ± 0.191 | ± 0.011 |
| 6 | -0.595 | -0.234 | 0.209 | -0.168 | -0.048 | 0.020 | 0.816 | 0.023 |
| | ± 0.268 | ± 0.269 | ± 0.150 | ± 0.162 | ± 0.154 | ± 0.102 | ± 0.209 | ± 0.010 |
| 1’’ | -0.048 | -0.511 | 0.071 | -0.050 | 0.146 | -0.047 | 0.683 | 0.022 |
| | ± 0.269 | ± 0.250 | ± 0.128 | ± 0.172 | ± 0.141 | ± 0.094 | ±0.190 | ± 0.010 |
| Single-Heme Reduced State | | | | | | | | |
| 6’ | -0.316 | -0.326 | 0.244 | -0.157 | 0.034 | 0.057 | 0.704 | 0.023 |
| | ± 0.254 | ± 0.279 | ± 0.152 | ± 0.161 | ± 0.164 | ± 0.108 | ± 0.166 | ± 0.012 |
| 1 | -0.097 | -0.429 | 0.047 | -0.100 | 0.157 | -0.039 | 0.623 | 0.023 |
| | ± 0.248 | ± 0.238 | ± 0.133 | ± 0.151 | ± 0.138 | ± 0.094 | ± 0.172 | ± 0.012 |
| 2 | -0.015 | -0.410 | 0.074 | -0.122 | 0.047 | 0.106 | 0.575 | 0.021 |
| | ± 0.220 | ± 0.209 | ± 0.120 | ± 0.145 | ± 0.140 | ± 0.088 | ± 0.161 | ± 0.009 |
| 3 | 0.029 | -1.134 | 0.051 | -0.123 | 0.030 | -0.072 | 1.187 | 0.021 |
| | ± 0.193 | ± 0.177 | ± 0.128 | ± 0.131 | ± 0.133 | ± 0.087 | ± 0.170 | ± 0.008 |
| 4 | -0.517 | -0.205 | 0.159 | -0.067 | -0.003 | -0.003 | 0.682 | 0.026 |
| | ± 0.249 | ± 0.204 | ± 0.130 | ± 0.143 | ± 0.133 | ± 0.079 | ± 0.196 | ± 0.056 |
| 5 | -0.271 | -0.577 | 0.096 | -0.037 | 0.252 | -0.080 | 0.780 | 0.026 |
| | ± 0.218 | ± 0.207 | ± 0.140 | ± 0.138 | ± 0.136 | ± 0.081 | ± 0.175 | ± 0.011 |
| 6 | -0.740 | -0.177 | 0.235 | -0.157 | -0.049 | 0.016 | 0.917 | 0.024 |
| | ± 0.272 | ± 0.267 | ± 0.143 | ± 0.150 | ± 0.174 | ± 0.103 | ± 0.219 | ± 0.011 |
| 1’’ | -0.211 | -0.501 | 0.023 | -0.020 | 0.200 | -0.098 | 0.701 | 0.022 |
| | ± 0.240 | ± 0.221 | ± 0.143 | ± 0.147 | ± 0.133 | ± 0.099 | ± 0.171 | ± 0.010 |

Table S5. In-plane normal coordinate structure decomposition for the hemes of OmcZ using the minimal basis set of vibrational modes.

| Heme # | B2g | B1g | Eux | Euy | A1g | A2g | SSD | Err |
|---|---|---|---|---|---|---|---|---|
| All Hemes Oxidized State | | | | | | | | |
| 7’ | 0.013 | 0.013 | 0.008 | 0.011 | 0.125 | 0.010 | 0.236 | 0.009 |
| | ± 0.095 | ± 0.114 | ± 0.073 | ± 0.081 | ± 0.090 | ± 0.042 | ± 0.065 | ± 0.003 |
| 1 | -0.049 | 0.022 | 0.063 | -0.000 | 0.133 | 0.007 | 0.247 | 0.010 |
| | ± 0.089 | ± 0.115 | ± 0.082 | ± 0.077 | ± 0.078 | ± 0.042 | ± 0.071 | ± 0.004 |
| 2 | -0.028 | 0.021 | -0.017 | -0.005 | 0.155 | 0.004 | 0.248 | 0.009 |
| | ± 0.098 | ± 0.105 | ± 0.076 | ± 0.079 | ± 0.078 | ± 0.040 | ± 0.065 | ± 0.003 |
| 3 | -0.003 | 0.023 | 0.005 | 0.002 | 0.182 | 0.014 | 0.267 | 0.009 |
| | ± 0.096 | ± 0.122 | ± 0.074 | ± 0.070 | ± 0.077 | ± 0.045 | ± 0.069 | ± 0.003 |
| 4 | -0.016 | -0.023 | 0.018 | 0.004 | 0.165 | 0.024 | 0.253 | 0.009 |
| | ± 0.101 | ± 0.109 | ± 0.065 | ± 0.077 | ± 0.086 | ± 0.040 | ± 0.074 | ± 0.003 |
| 5 | -0.010 | -0.006 | 0.021 | -0.023 | 0.153 | 0.018 | 0.241 | 0.009 |
| | ± 0.100 | ± 0.098 | ± 0.073 | ± 0.072 | ± 0.081 | ± 0.039 | ± 0.068 | ± 0.003 |
| 6 | 0.008 | -0.004 | 0.020 | -0.005 | 0.152 | 0.014 | 0.244 | 0.009 |
| | ± 0.104 | ± 0.107 | ± 0.072 | ± 0.071 | ± 0.080 | ± 0.043 | ± 0.070 | ± 0.003 |
| 7 | -0.001 | 0.001 | 0.030 | -0.003 | 0.115 | 0.007 | 0.238 | 0.010 |
| | ± 0.103 | ± 0.127 | ± 0.079 | ± 0.076 | ± 0.085 | ± 0.039 | ± 0.069 | ± 0.004 |
| 1’’ | -0.027 | 0.045 | 0.066 | -0.039 | 0.116 | 0.006 | 0.239 | 0.010 |
| | ± 0.095 | ± 0.107 | ± 0.072 | ± 0.075 | ± 0.085 | ± 0.041 | ± 0.073 | ± 0.004 |
| 8 | -0.011 | 0.006 | 0.015 | -0.009 | 0.178 | 0.013 | 0.264 | 0.009 |
| | ± 0.097 | ± 0.116 | ± 0.071 | ± 0.073 | ± 0.089 | ± 0.042 | ± 0.073 | ± 0.003 |
| Single-Heme Reduced State | | | | | | | | |
| 7’ | -0.019 | 0.031 | 0.013 | -0.011 | 0.182 | 0.012 | 0.260 | 0.008 |
| | ± 0.101 | ± 0.101 | ± 0.065 | ± 0.067 | ± 0.089 | ± 0.040 | ± 0.075 | ± 0.003 |
| 1 | -0.027 | 0.023 | 0.030 | -0.032 | 0.129 | 0.012 | 0.247 | 0.010 |
| | ± 0.107 | ± 0.119 | ± 0.080 | ± 0.073 | ± 0.090 | ± 0.044 | ± 0.077 | ± 0.003 |
| 2 | -0.018 | -0.008 | -0.014 | -0.003 | 0.188 | 0.008 | 0.270 | 0.009 |
| | ± 0.094 | ± 0.112 | ± 0.075 | ± 0.077 | ± 0.086 | ± 0.040 | ± 0.071 | ± 0.003 |
| 3 | 0.004 | 0.029 | 0.010 | 0.026 | 0.184 | 0.015 | 0.263 | 0.009 |
| | ± 0.096 | ± 0.110 | ± 0.066 | ± 0.072 | ± 0.082 | ± 0.040 | ± 0.075 | ± 0.003 |
| 4 | -0.017 | 0.004 | 0.008 | -0.040 | 0.171 | 0.007 | 0.251 | 0.009 |
| | ± 0.090 | ± 0.100 | ± 0.075 | ±0.066 | ± 0.079 | ±0.040 | ± 0.062 | ± 0.003 |
| 5 | -0.013 | -0.003 | 0.032 | -0.027 | 0.143 | 0.013 | 0.240 | 0.009 |
| | ± 0.092 | ± 0.105 | ± 0.070 | ± 0.076 | ± 0.090 | ± 0.042 | ± 0.070 | ± 0.003 |
| 6 | 0.018 | 0.007 | 0.030 | -0.031 | 0.113 | 0.017 | 0.230 | 0.010 |
| | ± 0.107 | ± 0.104 | ± 0.071 | ± 0.077 | ±0.091 | ± 0.038 | ± 0.068 | ± 0.003 |
| 7 | -0.019 | 0.004 | 0.012 | -0.006 | 0.172 | 0.009 | 0.257 | 0.008 |
| | ± 0.095 | ± 0.113 | ± 0.076 | ± 0.068 | ± 0.081 | ± 0.038 | ± 0.064 | ± 0.003 |
| 1’’ | -0.045 | 0.014 | 0.047 | -0.030 | 0.118 | 0.011 | 0.239 | 0.010 |
| | ± 0.092 | ± 0.107 | ± 0.075 | ± 0.081 | ± 0.098 | ± 0.041 | ± 0.072 | ± 0.004 |
| 8 | -0.013 | 0.030 | -0.007 | -0.019 | 0.171 | 0.013 | 0.256 | 0.008 |
| | ± 0.091 | ± 0.114 | ± 0.073 | ± 0.076 | ±0.080 | ± 0.040 | ± 0.073 | ± 0.003 |

Table S6. Out-plane normal coordinate structure decomposition for the hemes of OmcZ using the minimal basis set of vibrational modes.

| Heme # | B2u | B1u | A2u | Egx | Egy | A1u | SSD | Err |
|---|---|---|---|---|---|---|---|---|
| All Hemes Oxidized State | | | | | | | | |
| 7’ | -0.595 | -0.424 | -0.207 | -0.144 | 0.142 | 0.013 | 0.921 | 0.024 |
| | ± 0.368 | ± 0.284 | ± 0.156 | ± 0.178 | ± 0.166 | ± 0.105 | ±0.279 | ± 0.010 |
| 1 | -0.007 | -0.641 | 0.199 | -0.325 | -0.069 | -0.012 | 0.849 | 0.029 |
| | ± 0.243 | ± 0.280 | ± 0.134 | ± 0.167 | ± 0.133 | ± 0.090 | ± 0.220 | ± 0.015 |
| 2 | -0.044 | -0.233 | 0.074 | 0.215 | 0.314 | 0.022 | 0.608 | 0.028 |
| | ± 0.244 | ± 0.233 | ± 0.146 | ± 0.143 | ± 0.153 | ± 0.103 | ± 0.161 | ± 0.014 |
| 3 | -0.589 | 0.011 | 0.265 | -0.040 | 0.078 | 0.041 | 0.764 | 0.023 |
| | ± 0.285 | ± 0.243 | ± 0.135 | ± 0.143 | ± 0.150 | ± 0.096 | ± 0.229 | ± 0.011 |
| 4 | -0.661 | 0.075 | 0.020 | -0.044 | -0.255 | -0.003 | 0.823 | 0.022 |
| | ± 0.302 | ± 0.260 | ± 0.145 | ± 0.139 | ±0.152 | ± 0.096 | ± 0.251 | ± 0.010 |
| 5 | 0.295 | -0.446 | 0.063 | -0.123 | -0.001 | 0.056 | 0.701 | 0.023 |
| | ± 0.311 | ± 0.240 | ± 0.140 | ± 0.158 | ± 0.167 | ± 0.104 | ± 0.231 | ± 0.011 |
| 6 | -0.054 | -0.551 | -0.069 | -0.079 | 0.087 | 0.003 | 0.731 | 0.023± |
| | ± 0.340 | ± 0.254 | ± 0.142 | ± 0.162 | ± 0.156 | ± 0.097 | ± 0.224 | 0 .010 |
| 7 | -0.285 | -0.731 | -0.135 | -0.193 | 0.145 | -0.055 | 0.924 | 0.025 |
| | ± 0.265 | ± 0.262 | ± 0.137 | ± 0.142 | ± 0.142 | ± 0.092 | ± 0.215 | ± 0.010 |
| 1’’ | 0.385 | -0.611 | 0.117 | -0.435 | -0.047 | 0.018 | 0.936 | 0.028 |
| | ±0.267 | ± 0.219 | ± 0.133 | ± 0.140 | ± 0.150 | ± 0.089 | ± 0.190 | ± 0.012 |
| 8 | -0.293 | -0.269 | 0.052 | -0.142 | 0.169 | 0.054 | 0.670 | 0.023 |
| | ± 0.334 | ± 0.308 | ± 0.152 | ± 0.162 | ± 0.155 | ± 0.099 | ± 0.227 | ± 0.010 |
| Single-Heme Reduced State | | | | | | | | |
| 7’ | 0.072 | -0.328 | 0.036 | -0.234 | -0.004 | 0.018 | 0.594 | 0.025 |
| | ± 0.296 | ± 0.252 | ± 0.148 | ± 0.138 | ± 0.139 | ± 0.101 | ± 0.192 | ± 0.014 |
| 1 | 0.131 | -0.615 | 0.222 | -0.332 | -0.171 | 0.048 | 0.848 | 0.028 |
| | ± 0.236 | ± 0.213 | ± 0.130 | ± 0.151 | ± 0.139 | ± 0.099 | ± 0.190 | ± 0.012 |
| 2 | -0.111 | -0.226 | 0.092 | 0.112 | 0.187 | 0.003 | 0.536 | 0.025 |
| | ± 0.260 | ± 0.248 | ± 0.143 | ± 0.146 | ± 0.137 | ± 0.099 | ± 0.175 | ± 0.012 |
| 3 | -0.542 | -0.060 | 0.253 | -0.117 | 0.094 | -0.021 | 0.720 | 0.023 |
| | ± 0.236 | ± 0.234 | ± 0.128 | ± 0.130 | ± 0.141 | ± 0.088 | ± 0.190 | ± 0.011 |
| 4 | -0.390 | -0.145 | 0.032 | 0.224 | -0.145 | -0.025 | 0.631 | 0.027 |
| | ± 0.261 | ± 0.233 | ± 0.146 | ± 0.136 | ± 0.155 | ± 0.089 | ± 0.203 | ± 0.012 |
| 5 | 0.572 | -0.336 | 0.011 | -0.279 | 0.071 | 0.003 | 0.840 | 0.024 |
| | ± 0.306 | ±0.257 | ± 0.154 | ± 0.154 | ± 0.155 | ± 0.097 | ± 0.240 | ± 0.010 |
| 6 | 0.488 | -0.599 | 0.039 | -0.225 | -0.050 | -0.054 | 0.961 | 0.023 |
| | ± 0.402 | ± 0.293 | ± 0.156 | ± 0.175 | ± 0.172 | ± 0.110 | ± 0.271 | ±0.010 |
| 7 | -0.158 | -0.416 | -0.037 | -0.140 | 0.093 | -0.054 | 0.653 | 0.024 |
| | ± 0.301 | ± 0.243 | ± 0.146 | ± 0.165 | ± 0.137 | ± 0.094 | ± 0.169 | ± 0.010 |
| 1’’ | 0.163 | -0.728 | 0.203 | -0.261 | -0.068 | -0.045 | 0.910 | 0.024 |
| | ± 0.244 | ± 0.247 | ± 0.151 | ± 0.163 | ± 0.149 | ± 0.096 | ± 0.208 | ± 0.012 |
| 8 | -0.390 | -0.081 | 0.073 | -0.148 | 0.148 | 0.032 | 0.684 | 0.022 |
| | ± 0.350 | ± 0.312 | ± 0.148 | ± 0.172 | ± 0.155 | ± 0.097 | ± 0.209 | ± 0.011 |

Table S7. Assessment of the influence of the electrostatic environment on heme redox potentials in OmcE by three different methods

| Heme # | $\langle E^{\circ}_{vac}\rangle$ | $\langle E^{\circ}_{full}\rangle$ | $\langle E^{\circ}_{prot}\rangle$ | $\langle E^{\circ}_{solv}\rangle$ | $\langle E^{\circ}_{full}\rangle - \langle E^{\circ}_{vac}\rangle$ | $(\langle E^{\circ}_{prot}\rangle - \langle E^{\circ}_{vac}\rangle) + (\langle E^{\circ}_{solv}\rangle - \langle E^{\circ}_{vac}\rangle)$ | $\langle E_{ox}\rangle - \langle E_{red}$ |
|---|---|---|---|---|---|---|---|
| 1 | 0.544 ± 0.012 | -0.443 ± 0.028 | -17.148 ± 0.035 | 17.228 ± 0.041 | -0.987 ± 0.030 | -1.009 ± 0.056 | -1.182 ± 0.024 |
| 2 | 0.574 ± 0.012 | -0.405 ± 0.030 | -17.123 ± 0.045 | 17.230 ± 0.053 | -0.979 ± 0.032 | -1.041 ± 0.071 | -0.987 ± 0.024 |
| 3 | 0.493 ± 0.011 | -0.396 ± 0.029 | -19.181 ± 0.036 | 19.255 ± 0.042 | -0.889 ± 0.031 | -0.912 ± 0.057 | -0.963 ± 0.025 |
| 4 | 0.583 ± 0.012 | -0.298 ± 0.028 | -17.101 ± 0.038 | 17.379 ± 0.044 | -0.881 ± 0.030 | -0.889 ± 0.060 | -1.001 ± 0.023 |

Table S8. Assessment of the influence of the electrostatic environment on heme redox potentials in OmcS by three different methods

| Heme # | $\langle E^\circ_{vac}\rangle$ | $\langle E^\circ_{full}\rangle$ | $\langle E^\circ_{prot}\rangle$ | $\langle E^\circ_{solv}\rangle$ | $\langle E^\circ_{full}\rangle - \langle E^\circ_{vac}\rangle$ | $(\langle E^\circ_{prot}\rangle - \langle E^\circ_{vac}\rangle) + (\langle E^\circ_{solv}\rangle - \langle E^\circ_{vac}\rangle)$ | $\langle E_{ox}\rangle - \langle E_{red}$ |
|---|---|---|---|---|---|---|---|
| 1 | 0.594 | -0.109 | -11.610 | 12.025 | -0.703 | -0.773 | -0.614 |
| | ± 0.027 | ± 0.067 | ± 0.086 | ± 0.074 | ± 0.072 | ± 0.132 | ± 0.023 |
| 2 | 0.564 | -0.184 | -11.575 | 11.960 | -0.748 | -0.743 | -0.804 |
| | ± 0.026 | ± 0.055 | ± 0.077 | ± 0.100 | ± 0.061 | ± 0.138 | ± 0.024 |
| 3 | 0.525 | -0.126 | -11.694 | 12.080 | -0.651 | -0.664 | -0.567 |
| | ± 0.022 | ± 0.060 | ± 0.068 | ± 0.056 | ± 0.064 | ± 0.107 | ± 0.018 |
| 4 | 0.615 | -0.225 | -13.649 | 13.953 | -0.840 | -0.926 | -0.719 |
| | ± 0.028 | ± 0.061 | ± 0.049 | ± 0.065 | ± 0.067 | ± 0.102 | ± 0.019 |
| 5 | 0.533 | -0.185 | -14.047 | 14.331 | -0.718 | -0.782 | -1.101 |
| | ± 0.023 | ± 0.058 | ± 0.067 | ± 0.069 | ± 0.062 | ± 0.112 | ± 0.021 |
| 6 | 0.600 | -0.121 | -11.837 | 12.275 | -0.721 | -0.762 | -0.680 |
| | ± 0.021 | ± 0.052 | ± 0.058 | ± 0.068 | ± 0.056 | ± 0.103 | ± 0.021 |

Table S9. Assessment of the influence of the electrostatic environment on heme redox potentials in OmcZ by three different methods

| Heme # | $\langle E^{\circ}_{vac} \rangle$ | $\langle E^{\circ}_{full} \rangle$ | $\langle E^{\circ}_{prot} \rangle$ | $\langle E^{\circ}_{solv} \rangle$ | $\langle E^{\circ}_{full} \rangle - \langle E^{\circ}_{vac} \rangle$ | $(\langle E^{\circ}_{prot} \rangle - \langle E^{\circ}_{vac} \rangle) + (\langle E^{\circ}_{solv} \rangle - \langle E^{\circ}_{vac} \rangle)$ | $\langle E_{ox} \rangle - \langle E_{red}$ |
|---|---|---|---|---|---|---|---|
| 7’ | 0.555 ± 0.010 | -0.142 ± 0.029 | -17.326 ± 0.060 | 17.681 ± 0.075 | -0.697 ± 0.031 | -0.785 ± 0.102 | 0.735 ± 0.027 |
| 1 | 0.562 ± 0.010 | -0.331 ± 0.029 | -18.544 ± 0.056 | 18.739 ± 0.064 | -0.893 ± 0.030 | -0.921 ± 0.088 | -0.995 ± 0.022 |
| 2 | 0.536 ± 0.011 | -0.432 ± 0.027 | -19.534 ± 0.054 | 19.627 ± 0.067 | -0.968 ± 0.029 | -0.985 ± 0.086 | -1.056 ± 0.023 |
| 3 | 0.589 ± 0.010 | -0.451 ± 0.028 | -20.246 ± 0.044 | 20.384 ± 0.054 | -1.040 ± 0.029 | -1.056 ± 0.092 | -1.065 ± 0.024 |
| 4 | 0.551 ± 0.010 | -0.614 ± 0.029 | -19.694 ± 0.052 | 19.600 ± 0.059 | -1.165 ± 0.030 | -1.194 ± 0.080 | -1.295 ± 0.027 |
| 5 | 0.570 ± 0.011 | -0.570 ± 0.026 | -20.025 ± 0.065 | 19.980 ± 0.072 | -1.139 ± 0.028 | -1.168 ± 0.100 | -0.960 ± 0.029 |
| 6 | 0.574 ± 0.011 | -0.524 ± 0.028 | -19.038 ± 0.051 | 19.046 ± 0.065 | -1.097 ± 0.030 | -1.131 ± 0.089 | -1.291 ± 0.026 |
| 7 | 0.565 ± 0.011 | -0.266 ± 0.026 | -17.866 ± 0.052 | 18.136 ± 0.060 | -0.831 ± 0.028 | -0.872 ± 0.087 | -1.034 ± 0.027 |
| 1’’ | 0.549 ± 0.010 | -0.358 ± 0.026 | -18.222 ± 0.046 | 18.383 ± 0.059 | -0.907 ± 0.028 | -0.960 ± 0.079 | -1.019 ± 0.029 |
| 8 | 0.542 ± 0.011 | -0.580 ± 0.029 | -18.901 ± 0.066 | 18.877 ± 0.79 | -1.121 ± 0.031 | -1.126 ± 0.104 | -1.069 ± 0.022 |

Table S10. Cartesian coordinates for the lowed-energy gas-phase conformer found for the heme cofactor (not including propionic acid substituents) in the oxidized state using B3LYP/[Fe = LANL2DZ; H, C, N, S = 6-31G(d)]. The SCF energy was -2832.31734388 Hartrees.

| Atomic Symbol | X (Å) | Y (Å) | Z (Å) |
|---|---|---|---|
| Fe | 48.73859500 | 59.18494200 | 89.56658300 |
| N | 50.31669400 | 58.65836100 | 90.70486000 |
| C | 51.17071700 | 57.59273200 | 90.53680400 |
| C | 52.16187000 | 57.58341500 | 91.57934200 |
| C | 51.91404900 | 58.64916500 | 92.39470100 |
| C | 52.66372400 | 59.07455000 | 93.62021400 |
| H | 53.49548300 | 58.39315900 | 93.82015700 |
| H | 53.07915500 | 60.08415900 | 93.51192900 |
| H | 52.01946200 | 59.08315600 | 94.50820800 |
| C | 50.74946000 | 59.31466700 | 91.83198800 |
| C | 50.16837400 | 60.44932600 | 92.37853600 |
| H | 50.62400600 | 60.84580000 | 93.27842100 |
| C | 49.06121800 | 61.12383500 | 91.87985500 |
| N | 48.36106300 | 60.77963400 | 90.75090600 |
| C | 48.49110100 | 62.30604400 | 92.49845000 |
| C | 49.00231100 | 62.93153900 | 93.76431800 |
| H | 48.23439000 | 63.54529800 | 94.23946300 |
| H | 49.30212100 | 62.16953200 | 94.49155300 |
| H | 49.87910500 | 63.56615700 | 93.57800500 |
| C | 47.43350300 | 62.68571200 | 91.70714000 |
| C | 46.49128300 | 63.84850000 | 91.88782500 |
| H | 45.80220400 | 63.88407100 | 91.03831800 |
| C | 47.20096000 | 65.20929900 | 91.96652700 |
| H | 46.47460600 | 66.02494100 | 92.03448700 |
| H | 47.86587000 | 65.27556200 | 92.83054600 |
| H | 47.79772600 | 65.37031000 | 91.06017800 |
| C | 47.35986800 | 61.71384900 | 90.62852300 |
| C | 46.40657800 | 61.71143500 | 89.61702500 |
| H | 45.66312100 | 62.49830800 | 89.62407700 |
| C | 46.30971100 | 60.76957900 | 88.59842700 |
| N | 47.15983200 | 59.70315600 | 88.42353100 |
| C | 45.29163000 | 60.79244500 | 87.56934300 |
| C | 44.20219600 | 61.82048900 | 87.46903300 |
| H | 43.82874800 | 62.10791700 | 88.45797400 |
| H | 44.55172300 | 62.73684900 | 86.97444000 |
| H | 43.35343900 | 61.43254600 | 86.90211300 |
| C | 45.54302200 | 59.71520500 | 86.75158200 |
| C | 44.80984900 | 59.28221800 | 85.50812600 |
| H | 45.27823500 | 58.37862500 | 85.10864300 |
| C | 44.82777700 | 60.34088700 | 84.39166000 |
| H | 45.86272600 | 60.59770100 | 84.14025300 |

| | | | |
|---|---|---|---|
| H | 44.33865600 | 59.95540100 | 83.49169600 |
| H | 44.31474700 | 61.25864800 | 84.69261800 |
| C | 46.71155900 | 59.04455200 | 87.30479300 |
| C | 47.29478800 | 57.90019200 | 86.77310900 |
| H | 46.84848200 | 57.47838600 | 85.88099800 |
| C | 48.40481800 | 57.23566100 | 87.27406500 |
| N | 49.12087400 | 57.59634500 | 88.38976400 |
| C | 48.97086000 | 56.03970400 | 86.66981200 |
| C | 48.46372100 | 55.34963900 | 85.44024200 |
| H | 47.42753300 | 55.01003600 | 85.56187200 |
| H | 48.48725100 | 56.01030900 | 84.56463300 |
| H | 49.07497000 | 54.47250500 | 85.20986300 |
| C | 50.03736000 | 55.69887900 | 87.45018600 |
| C | 50.12657000 | 56.66532100 | 88.51162100 |
| C | 51.08864100 | 56.65424000 | 89.51501600 |
| H | 51.82827500 | 55.86010200 | 89.49750500 |
| C | 43.23033000 | 57.45124200 | 87.00318900 |
| H | 43.72335300 | 57.78550300 | 87.92067500 |
| H | 43.78894000 | 56.61917600 | 86.56309400 |
| S | 43.03219100 | 58.82231200 | 85.81463500 |
| C | 44.56744100 | 57.23825600 | 92.80704900 |
| H | 43.99045400 | 58.16545900 | 92.76730200 |
| H | 44.78323600 | 57.02578600 | 93.86123100 |
| C | 45.81802400 | 57.38713700 | 92.00530400 |
| N | 46.76307900 | 56.37749000 | 91.87535000 |
| H | 46.70995200 | 55.45944300 | 92.29532800 |
| C | 47.77488100 | 56.82065400 | 91.09758400 |
| H | 48.64298300 | 56.23624600 | 90.83805000 |
| N | 47.53469100 | 58.06610200 | 90.71547900 |
| C | 46.32078700 | 58.42891700 | 91.27282000 |
| H | 45.89697100 | 59.40719700 | 91.11631100 |
| C | 44.17719000 | 64.80382000 | 93.26261600 |
| H | 43.43653200 | 64.59145500 | 94.03834800 |
| H | 44.60602700 | 65.79101100 | 93.45249300 |
| S | 45.42820800 | 63.47555900 | 93.37188400 |
| C | 52.97948700 | 61.19803800 | 86.45420900 |
| H | 53.55986900 | 62.04413600 | 86.84205000 |
| H | 53.59352700 | 60.29972900 | 86.55559600 |
| C | 51.69597500 | 61.01927200 | 87.19558700 |
| N | 50.70781700 | 61.99391600 | 87.25011600 |
| H | 50.74777600 | 62.90206900 | 86.80788800 |
| C | 49.67767400 | 61.53303300 | 87.99312500 |
| H | 48.77711600 | 62.08989500 | 88.19656200 |
| N | 49.94632500 | 60.30963200 | 88.42324300 |
| C | 51.19789800 | 59.97946300 | 87.93404700 |
| H | 51.65159900 | 59.02508000 | 88.14487800 |

| | | | |
|---|---|---|---|
| H | 42.22264100 | 57.10466000 | 87.24738000 |
| H | 43.93102100 | 56.43065600 | 92.42492000 |
| H | 43.67447200 | 64.79653700 | 92.29014400 |
| H | 50.71331400 | 54.86247100 | 87.32008600 |
| H | 52.95123400 | 56.84738800 | 91.67209400 |
| H | 52.81173500 | 61.36817000 | 85.38366300 |

Table S11. Cartesian coordinates for the lowed-energy gas-phase conformer found for the heme cofactor (not including propionic acid substituents) in the reduced state using B3LYP/[Fe = LANL2DZ; H, C, N, S = 6-31G(d)]. The SCF energy was -2832.49574574 Hartrees.

| Atomic Symbol | X (Å) | Y (Å) | Z (Å) |
|---|---|---|---|
| Fe | 39.73142300 | 56.93291900 | 104.23919100 |
| N | 37.77134000 | 56.57719800 | 104.63021200 |
| C | 36.69217300 | 57.06792800 | 103.94416400 |
| C | 35.46284900 | 56.58146200 | 104.53256700 |
| C | 35.80114900 | 55.79104700 | 105.59049400 |
| C | 34.89294600 | 55.05472200 | 106.52931100 |
| H | 35.06785300 | 53.97078900 | 106.50428800 |
| H | 35.03348200 | 55.37471700 | 107.57027800 |
| H | 33.84330500 | 55.22581600 | 106.26911100 |
| C | 37.25990200 | 55.79706600 | 105.63501000 |
| C | 38.02179700 | 55.08302100 | 106.55681300 |
| H | 37.47747100 | 54.50449200 | 107.29658200 |
| C | 39.41486200 | 55.01615500 | 106.60088600 |
| N | 40.25872400 | 55.67172300 | 105.74469200 |
| C | 40.16365400 | 54.20519100 | 107.54840600 |
| C | 39.54635200 | 53.36710400 | 108.63120400 |
| H | 38.68908500 | 53.87505800 | 109.08835400 |
| H | 39.17980400 | 52.40424100 | 108.24740000 |
| H | 40.26583800 | 53.16106100 | 109.42737300 |
| C | 41.48998000 | 54.37367700 | 107.23448500 |
| C | 42.70217400 | 53.74182500 | 107.86270200 |
| H | 43.59905800 | 54.10050800 | 107.35052300 |
| C | 42.71105400 | 52.20637500 | 107.78196400 |
| H | 43.63612700 | 51.79934300 | 108.20513800 |
| H | 41.86549700 | 51.76764700 | 108.31941700 |
| H | 42.64360900 | 51.89536800 | 106.73355800 |
| C | 41.52631700 | 55.30656500 | 106.10725900 |
| C | 42.68620000 | 55.78737700 | 105.50016700 |
| H | 43.63319900 | 55.41966700 | 105.87841900 |
| C | 42.76147800 | 56.72008700 | 104.46599900 |
| N | 41.68749700 | 57.29557700 | 103.84321500 |
| C | 44.01120700 | 57.23422600 | 103.92547800 |
| C | 45.37922400 | 56.82115700 | 104.38747600 |
| H | 46.12636500 | 57.57542900 | 104.12917000 |
| H | 45.41181500 | 56.69066200 | 105.47559700 |
| H | 45.69103400 | 55.86584700 | 103.94204300 |
| C | 43.66532100 | 58.13849600 | 102.95152400 |
| C | 44.55493300 | 58.95195900 | 102.05134000 |
| H | 43.93327200 | 59.55834600 | 101.38689600 |
| C | 45.48361100 | 58.10510600 | 101.16563300 |
| H | 46.06750400 | 58.74242300 | 100.49232100 |

| | | | |
|---|---|---|---|
| H | 46.17907700 | 57.50772400 | 101.76191400 |
| H | 44.88363600 | 57.41757100 | 100.55916200 |
| C | 42.20298900 | 58.16156100 | 102.91794000 |
| C | 41.42930700 | 58.93162200 | 102.04816700 |
| H | 41.95145000 | 59.57942900 | 101.35207200 |
| C | 40.03911600 | 58.93334000 | 101.96029400 |
| N | 39.19777200 | 58.18716500 | 102.74277400 |
| C | 39.27971500 | 59.71844000 | 100.99158800 |
| C | 39.85308000 | 60.65344400 | 99.96920600 |
| H | 40.52863400 | 60.13710200 | 99.27418300 |
| H | 39.05752900 | 61.11420200 | 99.37467400 |
| H | 40.43204700 | 61.46333500 | 100.43268300 |
| C | 37.97063900 | 59.41379100 | 101.21743400 |
| C | 37.92910800 | 58.46469700 | 102.30904000 |
| C | 36.76082500 | 57.93816700 | 102.85743100 |
| H | 35.81972800 | 58.24982300 | 102.41169300 |
| C | 44.36604100 | 61.11269500 | 103.89075200 |
| H | 43.80471300 | 60.45124100 | 104.55622700 |
| H | 43.67306200 | 61.62778400 | 103.21802500 |
| S | 45.63102800 | 60.18233700 | 102.95956300 |
| C | 40.94761500 | 60.71792800 | 108.29194100 |
| H | 41.90410900 | 60.25344400 | 108.54696800 |
| H | 40.33524800 | 60.73617800 | 109.20326900 |
| C | 40.29380900 | 59.95110400 | 107.18983700 |
| N | 39.06732900 | 60.30166000 | 106.64472700 |
| H | 38.49668100 | 61.08717700 | 106.92039400 |
| C | 38.75888300 | 59.40254500 | 105.67070000 |
| H | 37.85667100 | 59.43769300 | 105.07992300 |
| N | 39.71245300 | 58.49896500 | 105.55942200 |
| C | 40.67106900 | 58.82891500 | 106.49702100 |
| H | 41.56604100 | 58.23646100 | 106.60790000 |
| C | 42.94187400 | 56.01936600 | 109.55208700 |
| H | 43.07020800 | 56.39595500 | 110.57075000 |
| H | 41.99002800 | 56.37942000 | 109.15199100 |
| S | 42.95546500 | 54.19723400 | 109.65879600 |
| C | 40.49359300 | 53.53654700 | 99.75754500 |
| H | 39.61953600 | 53.22950600 | 99.16796400 |
| H | 41.15224900 | 52.66321000 | 99.85425000 |
| C | 40.10701500 | 54.10024500 | 101.08516400 |
| N | 39.40995800 | 53.37831900 | 102.04316000 |
| H | 39.09365900 | 52.42398900 | 101.95490200 |
| C | 39.21867000 | 54.17867300 | 103.12689300 |
| H | 38.70097300 | 53.87018000 | 104.02186400 |
| N | 39.75094200 | 55.36730000 | 102.92114700 |
| C | 40.30540800 | 55.33075100 | 101.65720400 |
| H | 40.80773000 | 56.19266900 | 101.24628700 |

| | | | |
|---|---|---|---|
| H | 41.03165900 | 54.29274400 | 99.17935000 |
| H | 41.15067600 | 61.75908800 | 108.00808600 |
| H | 44.89850000 | 61.85833700 | 104.48785100 |
| H | 43.76507400 | 56.38919700 | 108.93244100 |
| H | 37.10175100 | 59.79723900 | 100.69446400 |
| H | 34.46695500 | 56.82188700 | 104.17760200 |

Table S12. Comparison of heme redox potentials in various homogenious solvents and the heterogenous OmcS protein nanowire[a]

| Solvent Environment | Heme # ϵ(298 K) | #1 | #2 | #3 | #4 | #5 | #6 |
|---|---|---|---|---|---|---|---|
| *n*-Hexane | 1.88 | 0.053 ± 0.024 | 0.047 ± 0.027 | 0.011 ± 0.026 | 0.094 ± 0.030 | 0.033 ± 0.025 | 0.100 ± 0.027 |
| Cyclohexane | 2.02 | 0.029 ± 0.029 | 0.028 ± 0.039 | -0.043 ± 0.037 | 0.055 ± 0.027 | -0.011 ± 0.023 | 0.065 ± 0.025 |
| *o*-Xylene | 2.35 | | -0.085 ± 0.030 | **-0.127 ± 0.024** | -0.028 ± 0.029 | -0.100 ± 0.027 | -0.044 ± 0.025 |
| Dibutylether | 3.05 | **-0.109 ± 0.030** | -0.139 ± 0.030 | -0.183 ± 0.025 | -0.087 ± 0.031 | -0.144 ± 0.027 | -0.079 ± 0.037 |
| Diethylamine | 3.58 | | **-0.169 ± 0.031** | | | **-0.185 ± 0.020** | **-0.120 ± 0.018** |
| Diethylether | 4.24 | -0.198 ± 0.027 | -0.212 ± 0.027 | -0.263 ± 0.023 | -0.152 ± 0.035 | -0.215 ± 0.028 | -0.145 ± 0.030 |
| Diethylsulfide | 5.72 | -0.225 ± 0.036 | -0.257 ± 0.030 | -0.311 ± 0.024 | -0.207 ± 0.033 | -0.262 ± 0.026 | -0.190 ± 0.031 |
| Ethanethiol | 6.67 | | | | **-0.228 ± 0.033** | | |
| 2,6-Dimethylpyridine | 7.17 | -0.254 ± 0.031 | -0.279 ± 0.030 | -0.347 ± 0.023 | -0.234 ± 0.029 | -0.285 ± 0.028 | -0.212 ± 0.031 |
| Tetrahydrofuran | 7.43 | -0.265 ± 0.029 | -0.285 ± 0.030 | -0.342 ± 0.025 | -0.243 ± 0.028 | -0.285 ± 0.028 | -0.218 ± 0.031 |
| OmcS | | -0.109 ± 0.067 | -0.184 ± 0.055 | -0.126 ± 0.060 | -0.225 ± 0.061 | -0.185 ±0.058 | -0.121 ± 0.052 |

[a]Redox potentials were computed for a common set of 26 conformers for each heme in each solvent environment. The homogeneous solvents were modeled with the polarizable continuum model, whereas snapshots from all-atom but non-polarizable MD was used for the protein context.

Table S13. Dependence of the electronic component of the heme redox potential on the solvent environment

| Environment | $U_{ox}$ (a.u.) | $U_{red}$ (a.u.) | E° |
|---|---|---|---|
| Vacuum | -2832.317344 | -2832.495746 | 0.536 |
| *n*-Hexane | -2832.344302 | -2832.503976 | 0.026 |
| Cyclohexane | -2832.346318 | -2832.504778 | -0.007 |
| *o*-Xylene | -2832.352252 | -2832.507322 | -0.099 |
| Dibutylether | -2832.356068 | -2832.509113 | -0.154 |
| Diethylamine | -2832.358998 | -2832.510577 | -0.194 |
| Diethylether | -2832.361700 | -2832.511999 | -0.229 |
| Diethylsulfide | -2832.365598 | -2832.514181 | -0.276 |
| Ethanethiol | -2832.367225 | -2832.515141 | -0.294 |
| 2,6-Dimethylpyridine | -2832.367932 | -2832.515568 | -0.302 |
| Tetrahydrofuran | -2832.368250 | -2832.515761 | -0.305 |
| Dichloroethane | -2832.370707 | -2832.517299 | -0.330 |
| Acetone | -2832.374280 | -2832.519673 | -0.363 |
| Methanol | -2832.375629 | -2832.520615 | -0.374 |
| Dimethylsulfoxide | -2832.376333 | -2832.521119 | -0.379 |
| Water | -2832.376992 | -2832.521596 | -0.384 |
| Formamide | -2832.377269 | -2832.521798 | -0.386 |
| *n*-Methylformamide-mixture | -2832.377554 | -2832.522008 | -0.388 |

Table S14. Vertical energy gaps and redox potentials for the hemes of OmcE with self-consistently optimized electron densities in vacuum and the protein environment, or with the density frozen at the vacuum-optimized distribution while the heme is in the protein context

| | Relaxed Density in Vacuum | | | Frozen Vacuum density In Protein Environment | | | Relaxed Density in Protein environment | | |
|---|---|---|---|---|---|---|---|---|---|
| Heme # | -VEA | VIP | E° | -VEA | VIP | E° | -VIA | VIP | E° |
| 4’ | 4.881 | 4.928 | 0.585 | 3.250 | 4.878 | -0.255 | 3.239 | 4.889 | -0.255 |
| | ± 0.008 | ± 0.009 | ± 0.012 | ± 0.019 | ± 0.018 | ± 0.026 | ± 0.018 | ± 0.019 | ± 0.02 |
| 1 | 4.874 | 4.852 | 0.544 | 2.961 | 4.742 | -0.467 | 3.013 | 4.739 | -0.443 |
| | ± 0.008 | ± 0.008 | ± 0.012 | ± 0.018 | ± 0.020 | ± 0.028 | ± 0.019 | ± 0.020 | ± 0.02 |
| 2 | 4.869 | 4.917± | 0.574 | 3.008 | 4.763 | -0.434 | 3.064 | 4.763 | -0.405 |
| | ± 0.007 | 0.009 | ± 0.012 | ± 0.021 | ± 0.019 | ± 0.028 | ± 0.023 | ± 0.020 | ± 0.03 |
| 3 | 4.808 | 4.816 | 0.493 | 3.084 | 4.705 | -0.424 | 3.135 | 4.710 | -0.396 |
| | ± 0.006 | ± 0.009 | ± 0.011 | ± 0.019 | ± 0.021 | ± 0.028 | ± 0.021 | ± 0.020 | ± 0.02 |
| 4 | 4.860 | 4.944 | 0.583 | 3.168 | 4.836 | -0.317 | 3.181 | 4.862 | -0.298 |
| | ± 0.007 | ± 0.009 | ± 0.012 | ± 0.021 | ± 0.040 | ± 0.045 | ± 0.021 | ± 0.018 | ± 0.02 |
| 1’’ | 4.882 | 4.863 | 0.554 | 2.970 | 4.871 | -0.398 | 3.044 | 4.865 | -0.365 |
| | ± 0.008 | ±0.008 | ± 0.012 | ± 0.024 | ± 0.023 | ± 0.033 | ± 0.024 | ± 0.021 | ± 0.03 |

Table S15. Vertical energy gaps and redox potentials for the hemes of OmcS with self-consistently optimized electron densities in vacuum and the protein environment, or with the density frozen at the vacuum-optimized distribution while the heme is in the protein context

| Heme # | Relaxed Density in Vacuum | | | Frozen Vacuum density In Protein Environment | | | Relaxed Density in Protein environment | | |
|---|---|---|---|---|---|---|---|---|---|
| | -VEA | VIP | E° | -VEA | VIP | E° | -VEA | VIP | E° |
| 6’ | 4.949 ± 0.008 | 4.930 ± 0.007 | 0.621 ± 0.011 | 3.449 ± 0.019 | 5.002 ± 0.017 | -0.093 ± 0.025 | 3.464 ± 0.020 | 5.003 ± 0.016 | -0.086 ± 0.026 |
| 1 | 4.888 ± 0.007 | 4.904 ± 0.008 | 0.577 ± 0.010 | 3.326 ± 0.021 | 5.027 ± 0.019 | -0.142 ± 0.028 | 3.372 ± 0.021 | 5.033 ± 0.020 | -0.117 ± 0.029 |
| 2 | 4.900 ± 0.007 | 4.912 ± 0.008 | 0.587 ± 0.011 | 3.327 ± 0.016 | 5.062 ± 0.018 | -0.125 ± 0.024 | 3.329 ± 0.016 | 5.047 ± 0.017 | -0.131 ± 0.023 |
| 3 | 4.834 ± 0.006 | 4.856 ± 0.006 | 0.526 ± 0.009 | 3.530 ± 0.015 | 4.825 ± 0.016 | -0.142 ± 0.022 | 3.555 ± 0.016 | 4.856 ± 0.015 | -0.114 ± 0.022 |
| 4 | 4.943 ± 0.008 | 4.932 ± 0.008 | 0.619 ± 0.011 | 3.344 ± 0.018 | 4.692 ± 0.018 | -0.301 ± 0.025 | 3.398 ± 0.019 | 4.691 ± 0.018 | -0.275 ± 0.026 |
| 5 | 4.870 ± 0.007 | 4.829 ± 0.007 | 0.530 ± 0.010 | 3.342 ± 0.019 | 4.889 ± 0.018 | -0.204 ± 0.026 | 3.333 ± 0.018 | 4.881 ± 0.018 | -0.212 ± 0.025 |
| 6 | 4.932 ± 0.008 | 4.935 ± 0.009 | 0.614 ± 0.012 | 3.392 ± 0.019 | 4.922 ± 0.018 | -0.162 ± 0.026 | 3.421 ± 0.019 | 4.938 ± 0.019 | -0.140 ± 0.027 |
| 1’’ | 4.869 ± 0.007 | 4.888 ± 0.007 | 0.559 ± 0.010 | 3.334 ± 0.019 | 5.027 ± 0.018 | -0.139 ± 0.026 | 3.356 ± 0.020 | 5.032 ± 0.018 | -0.125 ± 0.027 |

Table S16. Vertical energy gaps and redox potentials for the hemes of OmcZ with self-consistently optimized electron densities in vacuum and the protein environment, or with the density frozen at the vacuum-optimized distribution while the heme is in the protein context

| Heme # | Relaxed Density in Vacuum | | | Frozen Vacuum density In Protein Environment | | | Relaxed Density in Protein environment | | |
|---|---|---|---|---|---|---|---|---|---|
| | -VEA | VIP | E° | -VEA | VIP | E° | -VEA | VIP | E° |
| 7' | 4.869 ± 0.007 | 4.879 ± 0.007 | 0.555 ± 0.010 | 3.148 ± 0.021 | 5.126 ± 0.018 | -0.182 ± 0.028 | 3.207 ± 0.022 | 5.146 ± 0.018 | -0.142 ± 0.029 |
| 1 | 4.884 ± 0.008 | 4.877 ± 0.006 | 0.562 ± 0.010 | 3.147 ± 0.022 | 4.797 ± 0.016 | -0.347 ± 0.027 | 3.170 ± 0.024 | 4.806 ± 0.016 | -0.331 ± 0.029 |
| 2 | 4.848 ± 0.007 | 4.862 ± 0.008 | 0.536 ± 0.011 | 2.947 ± 0.019 | 4.815 ± 0.018 | -0.438 ± 0.026 | 2.953 ± 0.020 | 4.821 ± 0.019 | -0.432 ± 0.027 |
| 3 | 4.904 ± 0.007 | 4.912 ± 0.007 | 0.589 ± 0.010 | 2.811 ± 0.020 | 4.924 ± 0.019 | -0.452 ± 0.028 | 2.822 ± 0.021 | 4.913 ± 0.018 | -0.451 ± 0.028 |
| 4 | 4.884 ± 0.007 | 4.856 ± 0.007 | 0.551 ± 0.010 | 2.637 ± 0.019 | 4.721 ± 0.020 | -0.640 ± 0.028 | 2.682 ± 0.020 | 4.728 ± 0.020 | -0.614 ± 0.028 |
| 5 | 4.895 ± 0.008 | 4.883 ± 0.008 | 0.570 ± 0.011 | 2.783 ± 0.018 | 4.696 ± 0.019 | -0.580 ± 0.026 | 2.807 ± 0.018 | 4.692 ± 0.018 | -0.570 ± 0.026 |
| 6 | 4.894 ± 0.008 | 4.892 ± 0.008 | 0.574 ± 0.011 | 2.751 ± 0.020 | 4.824 ± 0.020 | -0.532 ± 0.028 | 2.770 ± 0.020 | 4.820 ± 0.019 | -0.524 ± 0.028 |
| 7 | 4.882 ± 0.008 | 4.885 ± 0.008 | 0.565 ± 0.011 | 3.067 ± 0.018 | 5.008 ± 0.019 | -0.281 ± 0.026 | 3.088 ± 0.018 | 5.019 ± 0.018 | -0.266 ± 0.026 |
| 1" | 4.866 ± 0.006 | 4.870 ± 0.008 | 0.549 ± 0.010 | 3.088 ± 0.017 | 4.810 ± 0.019 | -0.370 ± 0.026 | 3.108 ± 0.018 | 4.813 ± 0.018 | -0.358 ± 0.026 |
| 8 | 4.869 ± 0.007 | 4.853 ± 0.008 | 0.542 ± 0.011 | 2.563 ± 0.021 | 4.883 ± 0.022 | -0.596 ± 0.030 | 2.600 ± 0.020 | 4.879 ± 0.021 | -0.580 ± 0.029 |

Table S17. Decomposition of redox-linked changes in electrostatic interaction energy for each heme at 300 K in OmcE

| Heme # | 1 | 2 | 3 | 4 |
|---|---|---|---|---|
| Full Environment | -1.182 ± 0.024 | -0.987 ± 0.024 | -0.963 ± 0.025 | -1.001 ± 0.023 |
| Protein | -0.809 ± 0.027 | -0.458 ± 0.028 | -1.815 ± 0.030 | -0.856 ± 0.024 |
| Solvent | -0.373 ± 0.030 | -0.529 ± 0.032 | 0.852 ± 0.022 | -0.144 ± 0.026 |
| Non-Polar | -0.268 ± 0.011 | -0.091 ± 0.012 | -0.240 ± 0.008 | -0.179 ± 0.012 |
| Aromatic | 0.093 ± 0.005 | -0.111 ± 0.007 | -0.117 ± 0.009 | 0.015 ± 0.002 |
| Polar | 0.196 ± 0.009 | 0.103 ± 0.009 | 0.141 ± 0.009 | 0.527 ± 0.010 |
| Acidic | -0.134 ± 0.005 | -0.316 ± 0.008 | -1.109 ± 0.018 | -0.130 ± 0.003 |
| Basic | -0.273 ± 0.027 | 0.654 ± 0.029 | 0.292 ± 0.008 | 0.145 ± 0.015 |
| Other Hemes | 0.659 ± 0.006 | 0.581 ± 0.006 | 0.694 ± 0.004 | 0.383 ± 0.005 |
| Propionic acid groups | -1.082 ± 0.012 | -1.277 ± 0.020 | -1.476 ± 0.015 | -1.618 ± 0.017 |

Table S18. Decomposition of redox-linked changes in electrostatic interaction energy for each heme at 300 K in OmcS

| Heme # | 1 | 2 | 3 | 4 | 5 | 6 |
|---|---|---|---|---|---|---|
| Full Environment | -0.614 | -0.804 | -0.567 | -0.719 | -1.101 | -0.68 |
| | ± 0.023 | ± 0.024 | ± 0.018 | ± 0.019 | ± 0.021 | ± 0.021 |
| Protein | -0.742 | -0.732 | -0.592 | -1.133 | -1.548 | -0.641 |
| | ± 0.029 | ± 0.023 | ± 0.017 | ± 0.019 | ± 0.021 | ± 0.025 |
| Solvent | 0.129 | -0.073 | 0.025 | 0.414 | 0.447 | -0.039 |
| | ± 0.029 | ± 0.023 | ± 0.009 | ± 0.011 | ± 0.015 | ± 0.024 |
| Non-Polar | -0.152 | -0.343 | -0.165 | -0.278 | -0.297 | -0.596 |
| | ± 0.011 | ± 0.010 | ± 0.011 | ± 0.013 | ± 0.009 | ± 0.009 |
| Aromatic | 0.055 | 0.302 | 0.032 | -0.078 | -0.011 | 0.073 |
| | ± 0.002 | ± 0.010 | ± 0.005 | ± 0.003 | ± 0.008 | ± 0.005 |
| Polar | 0.276 | 0.034 | 0.210 | 0.072 | -0.299 | -0.013 |
| | ± 0.010 | ± 0.012 | ± 0.008 | ± 0.008 | ± 0.015 | ± 0.009 |
| Acidic | -1.157 | -0.517 | -0.189 | -0.173 | -0.325 | -0.054 |
| | ± 0.012 | ± 0.010 | ± 0.003 | ± 0.004 | ± 0.010 | ± 0.005 |
| Basic | 1.239 | 0.244 | 0.457 | 0.364 | 0.526 | 0.711 |
| | ± 0.024 | ± 0.010 | ± 0.011 | ± 0.009 | ± 0.008 | ± 0.018 |
| Other Hemes | 0.671 | 0.451 | 0.687 | 0.489 | 0.790 | 0.447 |
| | ± 0.004 | ± 0.004 | ± 0.004 | ± 0.004 | ± 0.004 | ± 0.005 |
| Propionic acid groups | -1.676 | -0.902 | -1.624 | -1.529 | -1.932 | -1.21 |
| | ± 0.019 | ± 0.010 | ± 0.014 | ± 0.011 | ± 0.010 | ± 0.019 |

Table S19. Decomposition of redox-linked changes in electrostatic interaction energy for each heme at 300 K in OmcZ

| Heme # | 1 | 2 | 3 | 4 | 5 | 6 | 7 | 8 |
|---|---|---|---|---|---|---|---|---|
| Full Environment | -0.995 | -1.056 | -1.065 | -1.295 | -0.960 | -1.291 | -1.034 | -1.019 |
| | + 0.022 | ± 0.023 | ± 0.024 | ± 0.027 | ± 0.029 | ± 0.026 | ± 0.027 | ± 0.029 |
| Protein | -0.632 | -1.211 | -0.855 | -0.980 | -2.257 | -1.169 | -0.518 | -1.239 |
| | ± 0.033 | ± 0.032 | ± 0.026 | ± 0.029 | ± 0.050 | ± 0.029 | ± 0.024 | ± 0.040 |
| Solvent | -0.364 | 0.155 | -0.210 | -0.315 | 1.297 | -0.122 | -0.516 | 0.220 |
| | ± 0.032 | ± 0.032 | ± 0.024 | ± 0.032 | ± 0.052 | ± 0.034 | ± 0.032 | ± 0.042 |
| Non-Polar | -0.134 | 0.004 | -0.009 | 0.073 | 0.305 | 0.365 | 0.007 | 0.090 |
| | ± 0.012 | ± 0.012 | ± 0.007 | ± 0.007 | ± 0.008 | ± 0.013 | ± 0.014 | ± 0.005 |
| Aromatic | 0.019 | 0.123 | -0.056 | -0.231 | 0.117 | -0.080 | -0.050 | 0.015 |
| | ± 0.005 | ± 0.004 | ± 0.005 | ± 0.008 | ± 0.003 | ± 0.003 | ± 0.005 | ± 0.001 |
| Polar | -0.014 | 0.212 | 0.114 ± | -0.038 | 0.185 | 0.248 | -0.207 | -0.333 |
| | ± 0.012 | ± 0.015 | 0.016 | ± 0.017 | ± 0.013 | ± 0.014 | ± 0.015 | ± 0.019 |
| Acidic | -0.263 | -0.628 | -0.482 | -0.606 | -0.340 | -0.231 | -0.021 | 0.079 |
| | ± 0.007 | ± 0.006 | ± 0.008 | ± 0.008 | ± 0.012 | ± 0.008 | ± 0.006 | ± 0.020 |
| Basic | 0.326 | 0.130 | 0.424 ± | 1.217 | 0.438 | 0.242 | 0.284 | 0.105 |
| | ± 0.019 | ± 0.022 | 0.010 | ± 0.015 | ± 0.011 | ± 0.011 | ± 0.010 | ± 0.010 |
| Other Hemes | 0.622 | 0.620 | 0.824 ± | 1.067 | 0.583 | 0.454 | 0.882 | 0.322 |
| | ± 0.005 | ± 0.005 | 0.006 | ± 0.006 | ± 0.008 | ± 0.007 | ± 0.006 | ± 0.003 |
| Propionic acid groups | -1.188 | -1.672 | -1.67 ± | -2.461 | -3.545 | -2.165 | -1.412 | -1.517 |
| | ± 0.032 | ± 0.023 | 0.032 | ± 0.023 | ± 0.052 | ± 0.003 | ± 0.015 | ± 0.025 |

Table S20. Per-residue contributions to the heme redox potentials in OmcE >|0.025| V determined from the electrostatic interaction energy for heme oxidation

| Heme #1 (ID=650) | | Heme #2 (ID=653) | | Heme #3 (ID=656) | | Heme #4 (ID=) | |
|---|---|---|---|---|---|---|---|
| PRN-652 | -0.435 | PRN-654 | -0.553 | PRN-658 | -0.876 | PRN-660 | -0.953 |
| LYS-637 | -0.388 | PRN-655 | -0.406 | ASP-352 | -0.600 | PRN-661 | -0.446 |
| PRN-651 | -0.379 | PRN-657 | -0.268 | PRN-657 | -0.381 | PRO-354 | -0.197 |
| PRN-672 | -0.274 | ALA-343 | -0.171 | GLU-342 | -0.183 | PRN-639 | -0.179 |
| PRO-269 | -0.195 | LYS-410 | -0.123 | PHE-419 | -0.158 | LYS-435 | -0.144 |
| LEU-270 | -0.110 | ASP-236 | -0.106 | PRO-418 | -0.151 | LEU-429 | -0.135 |
| HIO-235 | -0.105 | CYO-307 | -0.100 | PRN-654 | -0.133 | GLU-404 | -0.087 |
| ASP-413 | -0.082 | ASP-312 | -0.087 | GLU-404 | -0.123 | CYO-430 | -0.068 |
| LYS-234 | -0.077 | SER-304 | -0.080 | ASP-122 | -0.105 | THR-58 | -0.058 |
| ASP-236 | -0.075 | PHE-306 | -0.074 | SER-304 | -0.088 | GLN-51 | -0.048 |
| CYO-255 | -0.042 | MET-308 | -0.063 | PRO-354 | -0.084 | PRO-392 | -0.047 |
| THR-260 | -0.038 | PRN-658 | -0.056 | CYO-405 | -0.078 | ASP-352 | -0.045 |
| PRN-673 | -0.037 | ASP-337 | -0.053 | MET-308 | -0.064 | PRN-658 | -0.042 |
| HIO-412 | -0.034 | ILE-303 | -0.049 | ALA-343 | -0.056 | ALA-365 | -0.035 |
| ASN-571 | -0.028 | ASP-352 | -0.048 | HIO-353 | -0.049 | LEU-68 | -0.034 |
| CYO-632 | 0.025 | HIO-235 | -0.040 | THR-422 | -0.047 | ALA-366 | -0.029 |
| LEU-237 | 0.026 | THR-340 | -0.038 | PRN-655 | -0.045 | GLY-368 | -0.027 |
| VAL-411 | 0.027 | PRO-418 | -0.033 | ASP-416 | -0.039 | CYO-53 | 0.025 |
| HIO-636 | 0.032 | ILE-254 | -0.032 | ASN-344 | -0.038 | SER-285 | 0.026 |
| ILE-268 | 0.033 | LEU-305 | -0.030 | GLY-346 | -0.038 | ILE-52 | 0.027 |
| ASP-569 | 0.038 | GLY-274 | -0.029 | PRO-123 | -0.037 | THR-381 | 0.028 |
| LYS-410 | 0.040 | PHE-246 | -0.027 | ASN-351 | -0.035 | VAL-359 | 0.028 |
| ARG-421 | 0.047 | GLU-342 | -0.025 | ASP-413 | -0.034 | LEU-391 | 0.030 |
| PRN-654 | 0.051 | LEU-237 | 0.026 | PRN-652 | -0.032 | THR-384 | 0.031 |
| HIO-259 | 0.054 | ARG-421 | 0.028 | LEU-429 | 0.027 | ASP-367 | 0.032 |
| ASN-272 | 0.060 | TRP-271 | 0.031 | CYO-307 | 0.031 | ILE-364 | 0.037 |
| PHE-257 | 0.070 | HIO-409 | 0.031 | LYS-234 | 0.042 | ARG-421 | 0.037 |
| CYO-258 | 0.133 | GLY-313 | 0.035 | HIO-60 | 0.043 | LEU-370 | 0.047 |
| ARG-273 | 0.157 | LEU-345 | 0.037 | LEU-345 | 0.043 | HIO-57 | 0.047 |
| HEH-653 | 0.295 | THR-245 | 0.045 | LEU-420 | 0.053 | GLY-371 | 0.055 |
| HEH-671 | 0.356 | ILE-339 | 0.066 | ASN-61 | 0.061 | LEU-403 | 0.056 |
| | | VAL-411 | 0.081 | ASN-121 | 0.062 | GLY-432 | 0.063 |
| | | SER-309 | 0.087 | SER-407 | 0.079 | SER-286 | 0.071 |
| | | HIO-311 | 0.092 | HIO-409 | 0.093 | HIO-434 | 0.102 |
| | | CYO-310 | 0.117 | CYO-408 | 0.128 | HEH-656 | 0.105 |
| | | HEH-650 | 0.261 | ARG-421 | 0.144 | CYO-433 | 0.134 |

| | | | | | | | |
|---|---|---|---|---|---|---|---|
| | | LYS-234 | 0.266 | HEH-659 | 0.272 | HIO-353 | 0.143 |
| | | HEH-656 | 0.320 | HEH-653 | 0.409 | HEH-638 | 0.279 |
| | | ARG-273 | 0.410 | | | ASN-369 | 0.331 |

Table S21. Per-residue contributions to the heme redox potentials in OmcS >|0.025| V determined from the electrostatic interaction energy for heme oxidation

| Heme #1 (ID=1280) | | Heme #2 (ID=1274) | | Heme #3 (ID=1277) | | Heme #4 (ID=1271) | | Heme #5 (ID=1268) | | Heme #6 (ID=1283) | |
|---|---|---|---|---|---|---|---|---|---|---|---|
| PRN-1281 | -0.794 | PRN-1276 | -0.428 | PRN-1278 | -0.900 | PRN-1273 | -0.899 | PRN-1270 | -0.913 | PRN-1285 | -0.497 |
| PRN-1282 | -0.535 | PRN-1275 | -0.382 | PRN-1279 | -0.375 | PRN-1272 | -0.363 | PRN-1272 | -0.542 | PRN-1284 | -0.377 |
| ASP-145 | -0.374 | GLU-1164 | -0.286 | PRN-1275 | -0.303 | ALA-334 | -0.190 | PRN-1269 | -0.416 | PRN-1299 | -0.216 |
| PRN-1320 | -0.320 | GLY-5 | -0.263 | GLY-84 | -0.232 | PRN-1269 | -0.189 | SER-341 | -0.285 | CYO-400 | -0.136 |
| ASP-1221 | -0.300 | SER-46 | -0.156 | ILE-188 | -0.135 | PRO-129 | -0.184 | ASP-340 | -0.235 | MET-422 | -0.132 |
| MET-36 | -0.215 | ASP-145 | -0.155 | ASP-145 | -0.101 | CYO-239 | -0.124 | ASN-327 | -0.153 | PRO-258 | -0.125 |
| GLU-10 | -0.214 | TYR-62 | -0.149 | SER-44 | -0.084 | HIO-335 | -0.121 | CYO-328 | -0.130 | PRN-1270 | -0.108 |
| GLU-8 | -0.186 | GLN-43 | -0.103 | TYR-62 | -0.080 | MET-342 | -0.083 | ILE-246 | -0.125 | GLU-417 | -0.106 |
| LEU-37 | -0.107 | CYO-47 | -0.076 | ASP-85 | -0.075 | TRP-238 | -0.082 | PRO-258 | -0.111 | PHE-265 | -0.076 |
| ALA-7 | -0.097 | SER-44 | -0.071 | HIE-139 | -0.068 | PRO-134 | -0.067 | GLU-417 | -0.105 | PRO-295 | -0.071 |
| SER-46 | -0.092 | HIE-63 | -0.064 | CYO-140 | -0.056 | SER-236 | -0.053 | MET-342 | -0.083 | ALA-405 | -0.059 |
| GLU-1164 | -0.090 | PRO-146 | -0.060 | PRN-1273 | -0.056 | ARG-333 | -0.049 | HIO-243 | -0.067 | LYS-264 | -0.051 |
| HIO-1218 | -0.083 | PRN-1278 | -0.059 | ALA-336 | -0.047 | HIE-139 | -0.047 | PRO-129 | -0.060 | ASN-327 | -0.049 |
| PRO-35 | -0.081 | GLY-11 | -0.054 | HIO-51 | -0.026 | GLU-106 | -0.046 | SER-236 | -0.054 | MET-443 | -0.048 |
| TYR-1065 | -0.027 | ILE-188 | -0.050 | SER-141 | 0.026 | TYR-133 | -0.039 | HIE-247 | -0.053 | LEU-399 | -0.042 |
| PRN-1306 | -0.025 | GLN-39 | -0.043 | PRO-82 | 0.027 | PRN-1279 | -0.035 | HIO-257 | -0.043 | ALA-263 | -0.035 |
| THR-14 | -0.024 | ASP-1221 | -0.037 | TYR-194 | 0.027 | MET-235 | -0.035 | PRN-1273 | -0.034 | GLY-266 | -0.034 |
| HIO-144 | -0.024 | GLU-8 | -0.036 | ASN-241 | 0.027 | ASN-327 | -0.034 | ARG-333 | -0.027 | LEU-444 | -0.033 |
| CYO-1217 | 0.028 | GLN-52 | -0.033 | TYR-231 | 0.035 | SER-341 | -0.032 | GLY-260 | -0.027 | ALA-398 | -0.032 |
| ASN-49 | 0.038 | LYS-149 | -0.026 | CYO-47 | 0.039 | GLU-297 | -0.030 | PRO-295 | -0.025 | ALA-259 | -0.028 |
| SER-141 | 0.039 | SER-45 | -0.025 | GLY-109 | 0.041 | ASP-340 | -0.028 | GLY-427 | 0.027 | PHE-345 | 0.026 |
| PRO-196 | 0.041 | CYO-143 | 0.025 | LEU-138 | 0.047 | GLY-83 | -0.026 | GLU-237 | 0.028 | ARG-344 | 0.027 |
| ASN-1215 | 0.041 | ASN-25 | 0.025 | ILE-112 | 0.052 | GLY-130 | -0.026 | MET-431 | 0.033 | LEU-272 | 0.028 |
| LEU-38 | 0.047 | ILE-64 | 0.031 | LEU-189 | 0.060 | ASP-122 | -0.025 | ASN-424 | 0.033 | ILE-269 | 0.029 |
| CYO-1214 | 0.060 | CYO-140 | 0.033 | HIO-335 | 0.060 | CYO-331 | 0.026 | ARG-107 | 0.042 | HIO-420 | 0.045 |

| | | | | | | | | | | | |
|---|---|---|---|---|---|---|---|---|---|---|---|
| ARG-151 | 0.073 | HIO-144 | 0.033 | HIO-144 | 0.061 | PHE-205 | 0.027 | GLU-297 | 0.042 | CYO-416 | 0.047 |
| GLN-39 | 0.075 | SER-170 | 0.035 | PHE-86 | 0.061 | THR-127 | 0.036 | HIO-423 | 0.050 | ARG-256 | 0.057 |
| HIO-13 | 0.087 | ARG-151 | 0.036 | THR-81 | 0.062 | GLY-109 | 0.040 | THR-421 | 0.071 | LYS-604 | 0.066 |
| LYS-149 | 0.093 | ARG-187 | 0.042 | GLY-83 | 0.077 | ARG-344 | 0.043 | SER-330 | 0.099 | ASP-407 | 0.076 |
| PHE-1 | 0.094 | HIO-2 | 0.045 | SER-142 | 0.094 | ILE-246 | 0.055 | HIO-332 | 0.107 | TYR-273 | 0.078 |
| GLY-11 | 0.123 | SER-61 | 0.052 | ARG-333 | 0.133 | HIO-332 | 0.062 | MET-422 | 0.148 | VAL-326 | 0.081 |
| VAL-6 | 0.135 | SER-168 | 0.057 | CYO-143 | 0.140 | LEU-138 | 0.063 | CYO-331 | 0.184 | HEH-1268 | 0.145 |
| CYO-12 | 0.158 | SER-3 | 0.064 | ARG-187 | 0.279 | ASN-111 | 0.074 | ARG-344 | 0.188 | CYO-403 | 0.154 |
| ARG-1211 | 0.163 | GLY-4 | 0.071 | HEH-1271 | 0.305 | HIO-110 | 0.127 | HEH-1283 | 0.306 | HIO-257 | 0.170 |
| LYS-197 | 0.194 | ARG-1211 | 0.089 | HEH-1274 | 0.356 | HEH-1277 | 0.147 | ARG-256 | 0.328 | LYS-402 | 0.221 |
| HEH-1274 | 0.263 | HEH-1280 | 0.098 | | | ALA-128 | 0.152 | HEH-1271 | 0.475 | LYS-406 | 0.237 |
| HEH-1319 | 0.400 | HIO-51 | 0.099 | | | ASN-241 | 0.157 | | | HEH-1298 | 0.298 |
| LYS-1220 | 0.704 | CYO-50 | 0.132 | | | ARG-107 | 0.304 | | | | |
| | | ASN-49 | 0.152 | | | HEH-1268 | 0.339 | | | | |
| | | HEH-1277 | 0.347 | | | | | | | | |
| | | PHE-1 | 0.421 | | | | | | | | |

Table S22. Per-residue contributions to the heme redox potentials in OmcZ >|0.025| V determined from the electrostatic interaction energy for heme oxidation

| Heme #1 (ID=562) | | Heme #2 (ID=553) | | Heme #3 (ID=559) | | Heme #4 (ID=560) | | Heme #5 (ID=541) | | Heme #6 (ID=547) | | Heme #7 (ID=544) | | Heme #8 (ID=556) | |
|---|---|---|---|---|---|---|---|---|---|---|---|---|---|---|---|
| P564 | -0.617 | P560 | -0.804 | P561 | -0.612 | P552 | -1.045 | P548 | -1.521 | P546 | -1.460 | P546 | -0.417 | P558 | -0.897 |
| P563 | -0.411 | D529 | -0.471 | P560 | -0.584 | P561 | -0.590 | P543 | -0.717 | P548 | -0.305 | P545 | -0.390 | P557 | -0.568 |
| V414 | -0.179 | P554 | -0.392 | P554 | -0.326 | P551 | -0.443 | P552 | -0.637 | P549 | -0.301 | P845 | -0.373 | D471 | -0.324 |
| D529 | -0.167 | P555 | -0.351 | P558 | -0.214 | D388 | -0.331 | P542 | -0.419 | D365 | -0.142 | V319 | -0.286 | D460 | -0.161 |
| H523 | -0.166 | L496 | -0.143 | T463 | -0.171 | P543 | -0.294 | P549 | -0.214 | P545 | -0.130 | Q320 | -0.168 | Q486 | -0.137 |
| P263 | -0.149 | E437 | -0.125 | E437 | -0.136 | D297 | -0.171 | D307 | -0.201 | K305 | -0.087 | P846 | -0.157 | Q461 | -0.111 |
| V493 | -0.114 | Q500 | -0.074 | V489 | -0.124 | P542 | -0.159 | C308 | -0.094 | N368 | -0.078 | D365 | -0.151 | E437 | -0.094 |
| E488 | -0.097 | P551 | -0.072 | E488 | -0.108 | Y296 | -0.139 | E310 | -0.080 | Q375 | -0.078 | P549 | -0.100 | N477 | -0.057 |
| A411 | -0.093 | P561 | -0.067 | D398 | -0.102 | Y391 | -0.135 | D297 | -0.055 | D307 | -0.068 | C335 | -0.092 | I469 | -0.055 |
| N494 | -0.093 | S495 | -0.056 | D297 | -0.086 | T463 | -0.109 | T304 | -0.051 | F366 | -0.064 | F366 | -0.086 | C472 | -0.050 |
| N527 | -0.087 | T298 | -0.049 | C487 | -0.080 | D398 | -0.065 | Q375 | -0.047 | T345 | -0.049 | H322 | -0.080 | K387 | -0.041 |
| S412 | -0.075 | C436 | -0.042 | I484 | -0.064 | T304 | -0.064 | C392 | -0.034 | V319 | -0.044 | Q364 | -0.077 | P560 | -0.039 |
| C530 | -0.071 | V284 | -0.038 | L474 | -0.054 | C392 | -0.055 | G466 | -0.030 | C369 | -0.041 | S334 | -0.073 | S459 | -0.033 |
| N526 | -0.050 | V289 | -0.037 | F458 | -0.054 | G294 | -0.053 | N368 | -0.028 | C350 | -0.035 | L318 | -0.061 | T463 | -0.027 |
| G492 | -0.038 | H497 | -0.037 | Q486 | -0.053 | H464 | -0.040 | H379 | 0.024 | L325 | 0.027 | L346 | -0.049 | N473 | 0.027 |
| Q500 | -0.035 | G435 | -0.037 | P552 | -0.052 | I295 | -0.033 | C372 | 0.024 | T376 | 0.027 | H816 | -0.037 | H465 | 0.034 |
| M417 | -0.027 | P285 | -0.036 | D462 | -0.045 | S381 | -0.028 | F370 | 0.029 | P542 | 0.031 | C812 | -0.037 | G468 | 0.035 |
| P514 | 0.026 | C530 | -0.033 | P483 | -0.045 | P560 | 0.024 | F300 | 0.029 | H322 | 0.035 | V352 | -0.035 | T467 | 0.038 |
| I54 | 0.033 | I499 | -0.032 | Q461 | -0.041 | F300 | 0.028 | L303 | 0.030 | C311 | 0.044 | Q813 | -0.035 | ¤550 | 0.045 |
| R421 | 0.037 | G492 | -0.032 | G294 | -0.038 | P301 | 0.032 | Q321 | 0.034 | A324 | 0.046 | W817 | -0.034 | T470 | 0.049 |
| L496 | 0.039 | D297 | -0.032 | C472 | -0.032 | N399 | 0.037 | Y296 | 0.049 | P344 | 0.050 | T345 | -0.030 | L474 | 0.079 |
| Q531 | 0.040 | L519 | -0.030 | P555 | -0.027 | K387 | 0.037 | C369 | 0.054 | L309 | 0.060 | V342 | -0.026 | H476 | 0.116 |
| F81 | 0.042 | N494 | -0.028 | D402 | 0.026 | I469 | 0.040 | A306 | 0.056 | H373 | 0.061 | N368 | -0.025 | C475 | 0.139 |
| H497 | 0.046 | C487 | -0.028 | G466 | 0.027 | P554 | 0.043 | V313 | 0.067 | P377 | 0.088 | I336 | -0.025 | ¤559 | 0.274 |

| | | | | | | | | | | | | | | | |
|---|---|---|---|---|---|---|---|---|---|---|---|---|---|---|---|
| G435 | 0.053 | V493 | -0.027 | P400 | 0.036 | A383 | 0.043 | H378 | 0.076 | C308 | 0.128 | N371 | 0.035 | D462 | 0.508 |
| S410 | 0.055 | G522 | -0.026 | G435 | 0.036 | T298 | 0.044 | P377 | 0.076 | N371 | 0.139 | T348 | 0.039 | | |
| C53 | 0.057 | V489 | -0.025 | G394 | 0.037 | ¤556 | 0.048 | P301 | 0.080 | C372 | 0.140 | Y691 | 0.039 | | |
| Q434 | 0.071 | S459 | 0.024 | C436 | 0.037 | I397 | 0.059 | S314 | 0.100 | L318 | 0.151 | P343 | 0.044 | | |
| G532 | 0.079 | H523 | 0.025 | H393 | 0.038 | H379 | 0.076 | K299 | 0.167 | ¤541 | 0.178 | V696 | 0.051 | | |
| P416 | 0.089 | T463 | 0.027 | S419 | 0.044 | T376 | 0.081 | C311 | 0.168 | R367 | 0.195 | A333 | 0.055 | | |
| R85 | 0.089 | L293 | 0.028 | T467 | 0.047 | K390 | 0.084 | H312 | 0.184 | ¤544 | 0.283 | L325 | 0.059 | | |
| H534 | 0.100 | G525 | 0.028 | N477 | 0.052 | C389 | 0.100 | ¤547 | 0.210 | | | N809 | 0.066 | | |
| C533 | 0.152 | Q434 | 0.029 | H491 | 0.055 | H393 | 0.102 | ¤550 | 0.375 | | | P344 | 0.068 | | |
| R485 | 0.183 | P288 | 0.029 | T298 | 0.065 | H378 | 0.156 | | | | | I326 | 0.079 | | |
| ¤262 | 0.298 | A283 | 0.033 | K387 | 0.071 | ¤559 | 0.490 | | | | | H351 | 0.107 | | |
| ¤553 | 0.323 | K450 | 0.033 | H464 | 0.077 | ¤541 | 0.549 | | | | | D315 | 0.134 | | |
| | | N290 | 0.033 | I397 | 0.089 | K299 | 0.797 | | | | | G349 | 0.142 | | |
| | | G294 | 0.036 | C490 | 0.107 | | | | | | | C350 | 0.161 | | |
| | | W537 | 0.037 | P418 | 0.125 | | | | | | | R367 | 0.278 | | |
| | | K501 | 0.038 | P551 | 0.147 | | | | | | | ¤547 | 0.369 | | |
| | | G532 | 0.039 | K299 | 0.153 | | | | | | | ¤844 | 0.517 | | |
| | | W521 | 0.041 | N460 | 0.179 | | | | | | | | | | |
| | | S419 | 0.041 | ¤553 | 0.251 | | | | | | | | | | |
| | | P287 | 0.042 | ¤550 | 0.266 | | | | | | | | | | |
| | | P416 | 0.049 | ¤556 | 0.289 | | | | | | | | | | |
| | | H440 | 0.056 | | | | | | | | | | | | |
| | | A438 | 0.066 | | | | | | | | | | | | |
| | | F292 | 0.068 | | | | | | | | | | | | |
| | | ¤562 | 0.078 | | | | | | | | | | | | |
| | | P286 | 0.103 | | | | | | | | | | | | |
| | | N460 | 0.129 | | | | | | | | | | | | |
| | | C439 | 0.155 | | | | | | | | | | | | |
| | | ¤559 | 0.525 | | | | | | | | | | | | |

Table S23. Vertical energy gaps and redox potentials evaluated for Heme #3 as a function of switching off (zeroing-out) the atomic partial charges on specific residues for a common set of configurations of the trimeric OmcS assembly

| Residue | Vertical Energy Gap (eV) Oxidized Trajectory | Reduced Trajectory | Redox Potential (eV) | $\Delta E_{Env,j}^{QM,i}$ (eV)[a] |
|---|---|---|---|---|
| WT | 3.577 ± 0.03 | 4.794 ± 0.03 | -0.133 ± 0.05 | 0.000 |
| F86 | 3.532 ± 0.03 | 4.759 ± 0.03 | -0.174 ± 0.05 | 0.040 |
| E106 | 3.648 ± 0.03 | 4.830 ± 0.04 | -0.080 ± 0.05 | -0.053 |
| R107 | 3.494 ± 0.04 | 4.675 ± 0.04 | -0.235 ± 0.05 | 0.101 |
| A128 | 3.587 ± 0.04 | 4.750 ± 0.03 | -0.151 ± 0.05 | 0.017 |
| P129 | 3.582 ± 0.03 | 4.803 ± 0.04 | -0.127 ± 0.05 | -0.007 |
| Y133 | 3.587 ± 0.03 | 4.768 ± 0.04 | -0.142 ± 0.05 | 0.008 |
| S142 | 3.519 ± 0.03 | 4.672 ± 0.04 | -0.224 ± 0.05 | 0.090 |
| R187 | 2.997 ± 0.04 | 4.010 ± 0.03 | -0.816 ± 0.05 | 0.683 |
| W238 | 3.627 ± 0.03 | 4.863 ± 0.04 | -0.074 ± 0.05 | -0.059 |
| N241 | 3.492 ± 0.04 | 4.683 ± 0.04 | -0.232 ± 0.05 | 0.098 |
| R333 | 3.169 ± 0.04 | 4.364 ± 0.04 | -0.552 ± 0.05 | 0.419 |

[a] $\Delta E_{Env,i}^{QM,j}$ is computed as $\Delta E_{Env,j}^{QM,i} = \left\langle E_{Env,j_{on}}^{QM,i} \right\rangle - \left\langle E_{Env,j_{off}}^{QM,i} \right\rangle$, where $E_{Env,j_{on}}^{QM,i}$ is the redox potential of heme $i$ in the wild-type (WT) protein, and $E_{Env,j_{off}}^{QM,i}$ is the redox potential of heme $i$ in the chimeric system with the atomic partial charges on residue $j$ turned off.

Table S24. Average interaction energy between Heme #3 in the oxidized and reduced states with specific residues, and the change in the average interaction energy upon oxidation

| Residue | $\langle E_{\mathrm{Env,j}}^{MM,i_{ox}} \rangle$ | $\langle E_{Env,j}^{MM,i_{red}} \rangle$ | $E_{Env,j}^{MM,i}$ [a] |
|---|---|---|---|
| F86 | 0.038 | -0.033 | 0.071 |
| E106 | -0.003 | -0.001 | -0.002 |
| R107 | 0.007 | 0.004 | 0.003 |
| A128 | 0.003 | 0.001 | 0.002 |
| P129 | -0.004 | -0.004 | 0.000 |
| Y133 | 0.000 | 0.001 | -0.001 |
| S142 | -0.051 | -0.150 | 0.099 |
| R187 | 0.419 | 0.145 | 0.274 |
| W238 | -0.001 | 0.013 | -0.013 |
| N241 | -0.002 | -0.031 | 0.029 |
| R333 | 0.282 | 0.143 | 0.139 |

[a] $E_{Env,j}^{MM,i}$ was computed as $\Delta E_{Env,j}^{MM,i} = \left\langle E_{Env,j}^{MM,i_{ox}} \right\rangle - \left\langle E_{Env,j}^{MM,i_{red}} \right\rangle$, where $\langle \ldots \rangle$ denotes thermal averaging over the trajectory in which Heme $i$ was either oxidized or reduced.

Table S25. Vertical energy gaps and redox potentials evaluated for Heme #4 as a function of switching off (zeroing-out) the atomic partial charges on specific residues for a common set of configurations of the trimeric OmcS assembly

| Residue | Vertical Energy Gap (eV) Oxidized Trajectory | Reduced Trajectory | Redox Potential (eV) | $\Delta E^{QM,i}_{Env,j}$ (eV)[a] |
|---|---|---|---|---|
| WT | 3.484 ± 0.04 | 4.605 ± 0.03 | -0.274 ± 0.05 | 0.000 |
| F86 | 3.457 ± 0.04 | 4.587 ± 0.03 | -0.297 ± 0.05 | 0.023 |
| E106 | 3.661 ± 0.04 | 4.713 ± 0.03 | -0.132 ± 0.05 | -0.142 |
| R107 | 3.120 ± 0.04 | 4.253 ± 0.03 | -0.633 ± 0.05 | 0.358 |
| A128 | 3.315 ± 0.04 | 4.447 ± 0.03 | -0.438 ± 0.05 | 0.164 |
| P129 | 3.643 ± 0.04 | 4.779 ± 0.03 | -0.108 ± 0.04 | -0.166 |
| Y133 | 3.549 ± 0.04 | 4.681 ± 0.03 | -0.204 ± 0.05 | -0.070 |
| S142 | 3.482 ± 0.05 | 4.625 ± 0.03 | -0.266 ± 0.05 | -0.009 |
| R187 | 3.393 ± 0.04 | 4.501 ± 0.03 | -0.372 ± 0.05 | 0.098 |
| W238 | 3.623 ± 0.04 | 4.649 ± 0.03 | -0.183 ± 0.05 | -0.091 |
| N241 | 3.328 ± 0.04 | 4.439 ± 0.03 | -0.436 ± 0.05 | 0.161 |
| R333 | 3.317 ± 0.04 | 4.436 ± 0.03 | -0.443 ± 0.05 | 0.168 |

[a]$\Delta E^{QM,j}_{Env,i}$ is computed as $\Delta E^{QM,i}_{Env,j} = \left\langle E^{QM,i}_{Env,j_{on}} \right\rangle - \left\langle E^{QM,i}_{Env,j_{off}} \right\rangle$, where $E^{QM,i}_{Env,j_{on}}$ is the redox potential of heme $i$ in the wild-type (WT) protein, and $E^{QM,i}_{Env,j_{off}}$ is the redox potential of heme $i$ in the chimeric system with the atomic partial charges on residue $j$ turned off.

Table S26. Average interaction energy between Heme #4 in the oxidized and reduced states with specific residues, and the change in the average interaction energy upon oxidation

| Residue | $\langle E_{Env,j}^{MM,i_{ox}} \rangle$ | $\langle E_{Env,j}^{MM,i_{red}} \rangle$ | $E_{Env,j}^{MM,i}$ [a] |
|---:|---:|---:|---:|
| F86 | 0.023 | 0.000 | 0.023 |
| E106 | -0.087 | -0.036 | -0.051 |
| R107 | 0.558 | 0.284 | 0.275 |
| A128 | 0.226 | 0.062 | 0.164 |
| P129 | -0.573 | -0.369 | -0.204 |
| Y133 | -0.150 | -0.100 | -0.051 |
| S142 | 0.041 | 0.039 | 0.001 |
| R187 | 0.008 | 0.000 | 0.008 |
| W238 | -0.065 | 0.016 | -0.081 |
| N241 | -0.008 | -0.172 | 0.165 |
| R333 | -0.039 | 0.006 | -0.045 |

[a] $E_{Env,j}^{MM,i}$ was computed as $\Delta E_{Env,j}^{MM,i} = \left\langle E_{Env,j}^{MM,i_{ox}} \right\rangle - \left\langle E_{Env,j}^{MM,i_{red}} \right\rangle$, where $\langle ... \rangle$ denotes thermal averaging over the trajectory in which Heme $i$ was either oxidized or reduced.

Table S27. Vertical energy gaps and redox potentials evaluated for Heme #5 as a function of switching off (zeroing-out) the atomic partial charges on specific residues for a common set of configurations of the trimeric OmcS assembly

| Residue | Vertical Energy Gap (eV) Oxidized Trajectory | Reduced Trajectory | Redox Potential (eV) | $\Delta E_{Env,j}^{QM,i}$ (eV)[a] |
|---|---|---|---|---|
| WT | 3.367 ± 0.03 | 4.745 ± 0.03 | -0.263 ± 0.04 | 0.000 |
| F86 | 3.354 ± 0.03 | 4.733 ± 0.03 | -0.276 ± 0.04 | 0.012 |
| E106 | 3.390 ± 0.03 | 4.781 ± 0.03 | -0.233 ± 0.04 | -0.042 |
| R107 | 3.266 ± 0.03 | 4.560 ± 0.03 | -0.386 ± 0.04 | 0.153 |
| A128 | 3.316 ± 0.03 | 4.699 ± 0.03 | -0.311 ± 0.04 | -0.074 |
| P129 | 3.449 ± 0.03 | 4.792 ± 0.03 | -0.199 ± 0.04 | -0.113 |
| Y133 | 3.377 ± 0.03 | 4.742 ± 0.03 | -0.260 ± 0.04 | 0.061 |
| S142 | 3.364 ± 0.03 | 4.730 ± 0.03 | -0.272 ± 0.04 | 0.013 |
| R187 | 3.286 ± 0.03 | 4.656 ± 0.03 | -0.346 ± 0.04 | 0.074 |
| W238 | 3.374 ± 0.03 | 4.740 ± 0.03 | -0.262 ± 0.04 | -0.084 |
| N241 | 3.317 ± 0.03 | 4.689 ± 0.03 | -0.316 ± 0.04 | 0.055 |
| R333 | 3.218 ± 0.03 | 4.595 ± 0.03 | -0.412 ± 0.04 | 0.096 |

[a] $\Delta E_{Env,i}^{QM,j}$ is computed as $\Delta E_{Env,j}^{QM,i} = \left\langle E_{Env,j_{on}}^{QM,i} \right\rangle - \left\langle E_{Env,j_{off}}^{QM,i} \right\rangle$, where $E_{Env,j_{on}}^{QM,i}$ is the redox potential of heme $i$ in the wild-type (WT) protein, and $E_{Env,j_{off}}^{QM,i}$ is the redox potential of heme $i$ in the chimeric system with the atomic partial charges on residue $j$ turned off.

Table S28. Average interaction energy between Heme #5 in the oxidized and reduced states with specific residues, and the change in the average interaction energy upon oxidation

| Residue | $\langle E_{Env,j}^{MM,i_{ox}} \rangle$ | $\langle E_{Env,j}^{MM,i_{red}} \rangle$ | $E_{Env,j}^{MM,i}$ [a] |
|---:|---:|---:|---:|
| F86 | 0.000 | 0.000 | 0.000 |
| E106 | -0.013 | -0.006 | -0.006 |
| R107 | 0.081 | 0.050 | 0.030 |
| A128 | 0.029 | 0.006 | 0.023 |
| P129 | -0.036 | 0.015 | -0.052 |
| Y133 | 0.001 | 0.001 | -0.001 |
| S142 | 0.000 | 0.000 | 0.000 |
| R187 | 0.000 | 0.000 | 0.000 |
| W238 | -0.022 | -0.026 | 0.004 |
| N241 | 0.000 | -0.009 | 0.009 |
| R333 | 0.010 | 0.027 | -0.017 |

[a] $E_{Env,j}^{MM,i}$ was computed as $\Delta E_{Env,j}^{MM,i} = \left\langle E_{Env,j}^{MM,i_{ox}} \right\rangle - \left\langle E_{Env,j}^{MM,i_{red}} \right\rangle$, where $\langle \ldots \rangle$ denotes thermal averaging over the trajectory in which Heme $i$ was either oxidized or reduced.

Table S29. Redox potential shifts ($\Delta E_{Env,j}^{QM,i}$) and redox-associated changes in per-residue pairwise electrostatic energies ($\Delta E_{Env,j}^{MM,i}$)

| | Heme #3 | | Heme #4 | | Heme #5 | |
|---|---|---|---|---|---|---|
| Residue | $\Delta E_{Env,j}^{QM,i}$ | $\Delta E_{Env,j}^{MM,i}$ | $\Delta E_{Env,j}^{QM,i}$ | $\Delta E_{Env,j}^{MM,i}$ | $\Delta E_{Env,j}^{QM,i}$ | $\Delta E_{Env,j}^{MM,i}$ |
| F86 | 0.040 | 0.071 | 0.023 | 0.023 | 0.012 | 0.000 |
| E106 | -0.053 | -0.002 | -0.142 | -0.051 | -0.042 | -0.006 |
| R107 | 0.101 | 0.003 | 0.358 | 0.275 | 0.153 | 0.030 |
| A128 | 0.017 | 0.002 | 0.164 | 0.164 | -0.074 | 0.023 |
| P129 | -0.007 | 0.000 | -0.166 | -0.204 | -0.113 | -0.052 |
| Y133 | 0.008 | -0.001 | -0.070 | -0.051 | 0.061 | -0.001 |
| S142 | 0.090 | 0.099 | -0.009 | 0.001 | 0.013 | 0.000 |
| R187 | 0.683 | 0.274 | 0.098 | 0.008 | 0.074 | 0.000 |
| W238 | -0.059 | -0.013 | -0.091 | -0.081 | -0.084 | 0.004 |
| N241 | 0.098 | 0.029 | 0.161 | 0.165 | 0.055 | 0.009 |
| R333 | 0.419 | 0.139 | 0.168 | -0.045 | 0.096 | -0.017 |

Table S30. Characterization of inter-heme H-bonds in OmcZ (there are none in Omc- E or S)

| Acceptor | Donor | Occupancy (%) |
|---|---|---|
| All-Oxidized Hemes | | |
| PRN_561@O | HEH_550@ND11 | 0.87 |
| PRN_552@O | HEH_541@ND12 | 0.86 |
| PRN_560@O | HEH_553@ND11 | 0.79 |
| PRN_546@O | HEH_547@ND12 | 0.70 |
| PRN_548@O | HEH_541@ND11 | 0.54 |
| Heme #1 Reduced (All others oxidized) | | |
| PRN_560@O | HEH_553@ND11 | 0.83 |
| PRN_552@O | HEH_541@ND12 | 0.80 |
| PRN_561@O | HEH_550@ND11 | 0.76 |
| Heme #2 Reduced (All others oxidized) | | |
| PRN_551@O | HEH_559@ND12 | 0.72 |
| PRN_552@O | HEH_541@ND12 | 0.70 |
| PRN_561@O | HEH_550@ND11 | 0.60 |
| PRN_548@O | HEH_541@ND11 | 0.55 |
| PRN_560@O | HEH_553@ND11 | 0.55 |
| PRN_546@O | HEH_547@ND12 | 0.32 |
| Heme #3 Reduced (All others oxidized) | | |
| PRN_552@O | HEH_541@ND12 | 0.87 |
| PRN_561@O | HEH_550@ND11 | 0.84 |
| PRN_551@O | HEH_559@ND12 | 0.46 |
| PRN_560@O | HEH_553@ND11 | 0.43 |
| PRN_548@O | HEH_541@ND11 | 0.41 |
| Heme #4 Reduced (All others oxidized) | | |
| PRN_561@O | HEH_550@ND11 | 0.86 |
| PRN_552@O | HEH_541@ND12 | 0.80 |
| PRN_560@O | HEH_553@ND11 | 0.70 |
| PRN_548@O | HEH_541@ND11 | 0.49 |
| Heme #5 Reduced (All others oxidized) | | |
| PRN_552@O | HEH_541@ND12 | 0.87 |
| PRN_561@O | HEH_550@ND11 | 0.78 |
| PRN_560@O | HEH_553@ND11 | 0.47 |
| Heme #6 Reduced (All others oxidized) | | |
| PRN_552@O | HEH_541@ND12 | 0.84 |
| PRN_561@O | HEH_550@ND11 | 0.82 |
| PRN_551@O | HEH_559@ND12 | 0.52 |
| PRN_560@O | HEH_553@ND11 | 0.55 |

| | | |
|---|---|---|
| Heme #7 Reduced (All others oxidized) | | |
| PRN_552@O | HEH_541@ND12 | 0.86 |
| PRN_560@O | HEH_553@ND11 | 0.72 |
| PRN_561@O | HEH_550@ND11 | 0.71 |
| Heme #8 Reduced (All others oxidized) | | |
| PRN_546@O | HEH_547@ND12 | 0.75 |
| PRN_552@O | HEH_541@ND12 | 0.62 |
| PRN_560@O | HEH_553@ND11 | 0.47 |
| PRN_548@O | HEH_541@ND11 | 0.44 |
| PRN_561@O | HEH_550@ND11 | 0.38 |
| PRN_551@O | HEH_559@ND12 | 0.12 |

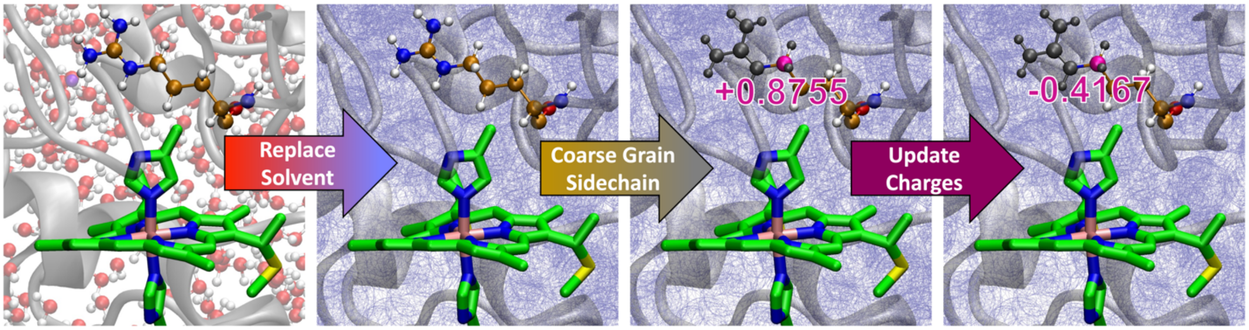


Figure S1. *In silico* mutagenesis strategy. First, the explicit solvent (water + counterions) is replaced by a bulk dielectric. This step has two advantages: It avoids the need to re-equilibrate the solvent in response to each considered mutation, and the polarization of the dielectric naturally handles the change in net charge for charge-changing mutations (i.e, no need to add/remove explicit ions). In the second step, the sidechain of the mutation-site is truncated (coarse-grained) to the last common atom between the original and final residue. This step avoids the introduction of bad steric clashes in the case of mutations that introduce larger groups. Finally, the mutation is modeled by updating the effective charge on the last common atom. Partial atomic charges on the other, shared atoms of the residue are also updated according to the force field definition of the new residue. This Effective Charge on Last Common Atom (ECLC) approach allows a rapid assessment of the influence of mutations on a quantum center because a fixed set of protein configurations is used, and only the electrostatic environment is perturbed. For ECLC to be valid, the assumption must be made—as is the case in any high-throughput computational protocol—that the mutation has a relatively small conformational effect.

Table S31. Marcus theory electron transfer energetic parameters for OmcE

| Heme Pair | $\Delta G°$ | $\lambda_{st}$ | $\lambda_{var,f}$ | $\lambda_{var,r}$ | $\lambda_{rxn}$ | $\langle H \rangle$ |
|---|---|---|---|---|---|---|
| 4'-1 | 0.188 | 0.710 | 0.724 | 0.741 | 0.688 | 7.797 |
| 1-2 | -0.038 | 0.931 | 1.044 | 0.928 | 0.879 | 1.472 |
| 2-3 | -0.009 | 0.587 | 0.764 | 0.666 | 0.481 | 13.001 |
| 3-4 | -0.099 | 0.766 | 0.747 | 0.756 | 0.780 | 1.775 |
| 4-1'' | 0.067 | 0.758 | 0.667 | 0.808 | 0.778 | 4.411 |

Table S32. Marcus theory electron transfer energetic parameters for OmcS

| Heme Pair | $\Delta G°$ | $\lambda_{st}$ | $\lambda_{var,f}$ | $\lambda_{var,r}$ | $\lambda_{rxn}$ | $\langle H \rangle$ |
|---|---|---|---|---|---|---|
| 6'-1 | 0.031 | 0.620 | 0.547 | 0.602 | 0.668 | 6.719 |
| 1-2 | 0.014 | 0.946 | 0.869 | 0.952 | 0.983 | 1.131 |
| 2-3 | -0.017 | 0.453 | 0.491 | 0.467 | 0.429 | 6.091 |
| 3-4 | 0.161 | 0.597 | 0.632 | 0.505 | 0.626 | 2.639 |
| 4-5 | -0.063 | 0.659 | 0.635 | 0.499 | 0.765 | 9.520 |
| 5-6 | -0.072 | 0.634 | 0.656 | 0.620 | 0.630 | 1.036 |
| 6-1'' | -0.015 | 0.676 | 0.625 | 0.639 | 0.724 | 7.895 |

Table S33. Marcus theory electron transfer energetic parameters for OmcZ

| Heme Pair | $\Delta G°$ | $\lambda_{st}$ | $\lambda_{var,f}$ | $\lambda_{var,r}$ | $\lambda_{rxn}$ | $\langle H \rangle$ |
|---|---|---|---|---|---|---|
| 7’-1 | 0.189 | 0.671 | 0.826 | 0.693 | 0.593 | 9.826 |
| 1-2 | 0.101 | 0.714 | 0.902 | 0.933 | 0.556 | 2.035 |
| 2-3 | 0.019 | 0.766 | 0.735 | 0.760 | 0.785 | 3.187 |
| 3-4 | 0.163 | 0.869 | 0.894 | 0.798 | 0.892 | 7.150 |
| 4-5 | -0.044 | 0.889 | 1.097 | 0.928 | 0.781 | 4.055 |
| 5-6 | -0.046 | 0.815 | 0.864 | 0.886 | 0.758 | 5.206 |
| 6-7 | -0.258 | 0.777 | 0.767 | 1.010 | 0.680 | 2.323 |
| 7-1’’ | 0.092 | 0.717 | 0.746 | 0.905 | 0.624 | 4.711 |

Table S34. Redox-state dependent solvent accessible surface areas for the heme cofactors

| Heme # | All-Hemes-Oxidized | Single-Heme-Reduced |
|---|---|---|
| | OmcE | |
| 4’ | 28.5 ± 9.0 | 14.0 ± 6.5 |
| 1 | 111.6 ± 15.1 | 81.2 ± 16.7 |
| 2 | 28.9 ± 8.0 | 15.8 ± 6.2 |
| 3 | 17.2 ± 4.7 | 24.6 ± 7.2 |
| 4 | 9.0 ± 11.6 | 4.5 ± 2.7 |
| 1’’ | 73.6 ± 17.8 | 88.9 ± 13.5 |
| | OmcS | |
| 6’ | 15.0 ± 4.6 | 16.3 ± 4.6 |
| 1 | 18.4 ± 4.9 | 22.7 ± 4.6 |
| 2 | 95.4 ± 8.2 | 63.9 ± 7.8 |
| 3 | 0.3 ± 0.4 | 0.3 ± 0.5 |
| 4 | 0.4 ± 0.6 | 0.6 ± 0.8 |
| 5 | 9.9 ± 3.8 | 2.7 ± 1.9 |
| 6 | 25.6 ± 5.4 | 13.3 ± 5.8 |
| 1’’ | 16.4 ± 5.8 | 27.1 ± 10.1 |
| | OmcZ | |
| 7’ | 27.9 ± 12.9 | 5.9 ± 3.2 |
| 1 | 12.1 ± 4.0 | 10.0 ± 4.4 |
| 2 | 16.0 ± 5.6 | 41.0 ± 12.3 |
| 3 | 11.6 ± 4.7 | 15.0 ± 6.3 |
| 4 | 72.5 ± 20.9 | 22.8 ± 15.6 |
| 5 | 147.1 ± 21.1 | 98.3 ± 16.3 |
| 6 | 176.3 ± 25.1 | 52.4 ± 14.8 |
| 7 | 31.2 ± 10.0 | 8.7 ± 4.7 |
| 1’’ | 22.9 + 5.8 | 18.2 ± 5.7 |

Table S35. Minimum distance (Å) between adjacent hemes in OmcE from molecular dynamics simulations in different redox microstates

| | All Oxidized | $4'_{\mathrm{red}}$ | $1_{\mathrm{red}}$ | $2_{\mathrm{red}}$ | $3_{\mathrm{red}}$ | $4_{\mathrm{red}}$ | $1'_{\mathrm{red}}$ |
|---|---|---|---|---|---|---|---|
| 4'-1 (S) | 4.5±0.3 | 4.0±0.2 | 4.4±0.3 | 4.4±0.4 | 4.2±0.3 | 4.2±0.3 | 4.4±0.3 |
| 1-2 (T) | 6.0±0.2 | 5.9±0.2 | 5.9± 0.2 | 5.9±0.2 | 5.9±0.2 | 6.1±0.2 | 6.1±0.2 |
| 2-3 (S) | 3.7±0.2 | 4.1±0.2 | 3.9 ±0.2 | 3.7±0.2 | 4.1±0.2 | 4.1±0.2 | 3.7±0.2 |
| 3-4 (T) | 6.0±0.2 | 5.9±0.2 | 6.0 ±0.2 | 6.0±0.2 | 5.9±0.2 | 5.9±0.2 | 6.0±0.2 |
| 4-1'' (S) | 4.4±0.3 | 3.9±0.2 | 4.2 ±0.3 | 4.2±0.3 | 4.0±0.2 | 3.9±0.2 | 4.3±0.3 |

Table S36. Minimum distance (in Å) between adjacent hemes in OmcS from molecular dynamics simulations in different redox microstates

| Heme Pair | All Ox-idized | $6'_{\mathrm{red}}$ | $1_{\mathrm{red}}$ | $2_{\mathrm{red}}$ | $3_{\mathrm{red}}$ | $4_{\mathrm{red}}$ | $5_{\mathrm{red}}$ | $6_{\mathrm{red}}$ | $1''_{red}$ |
|---|---|---|---|---|---|---|---|---|---|
| 6'-1 (S) | 4.1 ± 0.2 | 4.0 ± 0.2 | 4.1 ± 0.2 | 4.2 ± 0.3 | 4.3 ± 0.3 | 4.1 ± 0.2 | 4.0 ± 0.2 | 4.0 ± 0.2 | 4.2 ± 0.3 |
| 1-2 (T) | 6.1 ± 0.2 | 5.9 ± 0.2 | 5.9 ± 0.2 | 6.0 ± 0.2 | 6.0 ± 0.2 | 6.2 ± 0.2 | 6.0 ± 0.2 | 6.2 ± 0.3 | 6.2 ± 0.2 |
| 2-3 (S) | 4.1 ± 0.2 | 3.8 ± 0.2 | 4.0 ± 0.2 | 3.9 ± 0.2 | 3.9 ± 0.2 | 3.9 ± 0.2 | 4.0 ± 0.2 | 4.0 ± 0.2 | 3.9 ± 0.2 |
| 3-4 (T) | 6.0 ± 0.2 | 6.0 ± 0.2 | 6.0 ± 0.2 | 6.0 ± 0.2 | 6.0 ± 0.2 | 6.0 ± 0.2 | 6.0 ± 0.2 | 6.0 ± 0.2 | 6.1 ± 0.2 |
| 4-5 (S) | 3.7 ± 0.2 | 3.9 ± 0.2 | 4.1 ± 0.2 | 3.8 ± 0.2 | 3.9 ± 0.2 | 3.8 ± 0.2 | 3.8 ± 0.2 | 3.8 ± 0.2 | 3.9 ± 0.2 |
| 5-6 (T) | 6.0 ± 0.2 | 5.9 ± 0.2 | 5.9 ± 0.2 | 6.0 ± 0.2 | 5.9 ± 0.2 | 5.8 ± 0.2 | 5.9 ± 0.2 | 6.0 ± 0.2 | 6.0 ± 0.2 |
| 6-1'' (S) | 4.2 ± 0.3 | 4.1 ± 0.3 | 4.1 ± 0.3 | 4.1 ± 0.3 | 4.0 ± 0.2 | 4.3 ± 0.3 | 4.0 ± 0.2 | 4.0 ± 0.2 | 4.0 ± 0.2 |

Table S37. Minimum distance (in Å) between adjacent hemes along the main chain of OmcZ from molecular dynamics simulations in different redox microstates

| Heme Pair | All Ox-idized | $7'_{\mathrm{red}}$ | $1_{\mathrm{red}}$ | $2_{\mathrm{red}}$ | $3_{\mathrm{red}}$ | $4_{\mathrm{red}}$ | $5_{red}$ | $6_{red}$ | $7_{red}$ | $1''_{red}$ |
|---|---|---|---|---|---|---|---|---|---|---|
| 7'-1 (S) | 4.0 ± 0.3 | 3.7 ± 0.2 | 3.8 ± 0.2 | 4.1 ± 0.2 | 4.0 ± 0.3 | 3.9 ± 0.2 | 3.9 ± 0.2 | 4.3 ± 0.3 | 4.1 ± 0.3 | 3.9 ± 0.3 |
| 1-2 (T) | 5.5 ± 0.2 | 5.6 ± 0.2 | 5.4 ± 0.2 | 5.5 ± 0.2 | 5.4 ± 0.2 | 5.5 ± 0.2 | 5.7 ± 0.2 | 5.6 ± 0.2 | 5.6 ± 0.2 | 5.5 ± 0.2 |
| 2-3 (S) | 4.0 ± 0.2 | 3.9 ± 0.2 | 4.3 ± 0.2 | 4.1 ± 0.2 | 4.2 ± 0.2 | 4.0 ± 0.2 | 3.9 ± 0.2 | 3.9 ± 0.2 | 4.0 ± 0.2 | 3.9 ± 0.2 |
| 3-4 (S) | 4.2 ± 0.2 | 4.5 ± 0.3 | 4.0 ± 0.2 | 4.3 ± 0.2 | 4.2 ± 0.2 | 4.2 ± 0.2 | 4.8 ± 0.3 | 4.2 ± 0.2 | 4.0 ± 0.2 | 4.2 ± 0.3 |
| 4-5 (T) | 4.9 ± 0.2 | 4.8 ± 0.2 | 5.0 ± 0.3 | 5.0 ± 0.3 | 5.2 ± 0.3 | 5.3 ± 0.3 | 5.0 ± 0.3 | 4.9 ± 0.3 | 5.5 ± 0.3 | 5.0 ± 0.3 |
| 5-6 (S) | 4.0 ± 0.3 | 3.9 ± 0.3 | 3.7 ± 0.2 | 4.2 ± 0.3 | 4.2 ± 0.3 | 4.2 ± 0.2 | 3.7 ± 0.2 | 3.7 ± 0.2 | 3.8 ± 0.3 | 3.9 ± 0.3 |
| 6-7 (T) | 5.3 ± 0.3 | 5.1 ± 0.3 | 4.9 ± 0.3 | 5.1 ± 0.3 | 4.9 ± 0.2 | 5.0 ± 0.3 | 5.1 ± 0.3 | 4.9 ± 0.2 | 4.8 ± 0.2 | 5.1 ± 0.3 |
| 7-1'' (S) | 3.9 ± 0.2 | 4.0 ± 0.3 | 3.9 ± 0.3 | 4.3 ± 0.3 | 3.8 ± 0.2 | 4.2 ± 0.3 | 4.2 ± 0.3 | 4.0 ± 0.2 | 3.9 ± 0.2 | 4.0 ± 0.2 |

Table S38. Computed charge diffusion constants and redox currents[a]

| | Charge Diffusion Constant ($cm^2/s$) | Current (pA) at 0.1 V For a 300 nm filament |
|---|---|---|
| | Computed Energetic Parameters | |
| OmcE | $1.1 \times 10^{-8}$ / $4.7 \times 10^{-9}$ | $1.2 \times 10^{-3}$ / $5.4 \times 10^{-4}$ |
| OmcS | $7.8 \times 10^{-9}$ / $3.7 \times 10^{-9}$ | $9.8 \times 10^{-4}$ / $4.6 \times 10^{-4}$ |
| OmcZ | $1.3 \times 10^{-10}$ / $1.0 \times 10^{-10}$ | $1.7 \times 10^{-5}$ / $1.3 \times 10^{-5}$ |
| | $\Delta G = 0$ | |
| OmcE | $9.5 \times 10^{-9}$ / $1.3 \times 10^{-9}$ | $1.1 \times 10^{-3}$ / $1.5 \times 10^{-4}$ |
| OmcS | $9.9 \times 10^{-9}$ / $8.1 \times 10^{-9}$ | $1.3 \times 10^{-3}$ / $1.0 \times 10^{-3}$ |
| OmcZ | $9.2 \times 10^{-8}$ / $2.9 \times 10^{-7}$ | $1.2 \times 10^{-2}$ / $3.8 \times 10^{-2}$ |
| | $\Delta G = -\lambda_{rxn}$ | |
| OmcE | $2.2 \times 10^{-4}$ / $2.0 \times 10^{-4}$ | $2.7 \times 10^{1}$ / $2.4 \times 10^{1}$ |
| OmcS | $1.9 \times 10^{-4}$ / $1.9 \times 10^{-4}$ | $2.4 \times 10^{1}$ / $2.4 \times 10^{1}$ |
| OmcZ | $4.6 \times 10^{-4}$ / $4.0 \times 10^{-4}$ | $6.0 \times 10^{1}$ / $5.3 \times 10^{1}$ |

aValues given as X/Y indicate the result obtained by including the electron transfer step either from the last heme in the preceding subunit or to the first heme in the proceeding subunit.

Table S39. Reorganization energies computed for electron transfers in OmcZ in explicit methanol and chloroform solutions to model the low dielectric of the aggregated filament

| | Chloroform + Neutralizing $Na^+$ ions | | | | Neutralizing $Na^+$ ions | | | |
|---|---|---|---|---|---|---|---|---|
| Heme Pair | $\lambda_{st}$ | $\lambda_{var,f}$ | $\lambda_{var,b}$ | $\lambda_{rxn}$ | $\lambda_{st}$ | $\lambda_{var,f}$ | $\lambda_{var,b}$ | $\lambda_{rxn}$ |
| 7'-1 | 0.688 | 0.504 | 0.828 | 0.710 | 0.629 | 0.441 | 1.050 | 0.530 |
| 1-2 | 0.704 | 0.591 | 0.632 | 0.811 | 0.698 | 0.578 | 0.553 | 0.862 |
| 2-3 | 0.630 | 0.494 | 0.533 | 0.773 | 0.631 | 0.501 | 0.551 | 0.757 |
| 3-4 | 0.619 | 0.486 | 0.649 | 0.674 | 0.562 | 0.467 | 0.612 | 0.586 |
| 4-5 | 0.330 | 0.553 | 0.645 | 0.182 | 0.139 | 0.583 | 0.656 | 0.031 |
| 5-6 | 0.460 | 0.763 | 0.970 | 0.244 | 0.381 | 0.686 | 1.118 | 0.161 |
| 6-7 | 0.804 | 0.778 | 0.504 | 1.009 | 0.773 | 0.746 | 0.479 | 0.975 |
| 7-1" | 0.573 | 0.368 | 0.644 | 0.649 | 0.589 | 0.378 | 0.685 | 0.652 |
| | Methanol + Neutralizing $Na^+$ ions | | | | Neutralizing $Na^+$ ions | | | |
| | $\lambda_{st}$ | $\lambda_{var,f}$ | $\lambda_{var,b}$ | $\lambda_{rxn}$ | $\lambda_{st}$ | $\lambda_{var,f}$ | $\lambda_{var,b}$ | $\lambda_{rxn}$ |
| 7'-1 | 0.871 | 0.837 | 0.829 | 0.911 | 0.554 | 2.722 | 0.826 | 0.173 |
| 1-2 | 0.919 | 0.823 | 1.041 | 0.907 | 0.816 | 1.064 | 1.896 | 0.450 |
| 2-3 | 0.702 | 0.787 | 1.849 | 0.373 | 0.298 | 1.587 | 1.809 | 0.052 |
| 3-4 | 1.043 | 1.303 | 0.903 | 0.986 | 0.726 | 1.198 | 1.210 | 0.438 |
| 4-5 | 1.064 | 0.908 | 0.843 | 1.292 | 0.885 | 3.843 | 1.383 | 0.300 |
| 5-6 | 0.875 | 0.759 | 1.252 | 0.761 | 0.153 | 1.945 | 4.036 | 0.008 |
| 6-7 | 0.937 | 0.765 | 1.878 | 0.665 | 0.634 | 1.460 | 2.079 | 0.227 |
| 7-1" | 0.778 | 0.980 | 0.823 | 0.672 | 0.453 | 1.627 | 1.030 | 0.154 |

Table S40. Length (in ns) of production-stage molecular dynamics simulations performed in various redox microstates

| | OmcE | OmcS | OmcZ |
|---|---|---|---|
| All Oxidized | 144 | 288 | 108 |
| i = 1 | 416 | 301 | 176 / 180 / 180 |
| i = 2 | 416 | 281 | 176 / 180 / 180 |
| i = 3 | 416 | 266 | 176 / 180 / 174 |
| i = 4 | 416 | 295 | 176 / 180 / 173 |
| i = 5 | | 242 | 176 / 180 / 173 |
| i = 6 | | 294 | 176 / 216 / 173 |
| i = 7 | | | 176 / 180 / 173 |
| i = 8 | | | 142 / 180 / 173 |
| i - 1 | 416 | 72 | 162 / 180 / 180 |
| i + 1 | 216 | 72 | 148 / 180 / 173 |
